# A split ribozyme that links detection of a native RNA to orthogonal protein outputs

**DOI:** 10.1101/2022.01.12.476080

**Authors:** Lauren Gambill, August Staubus, Andrea Ameruoso, James Chappell

## Abstract

Individual RNA remains a challenging signal to synthetically transduce into different types of cellular information. Here, we describe Ribozyme-ENabled Detection of RNA (RENDR), a plug-and-play strategy that uses cellular transcripts to template the assembly of split ribozymes, triggering splicing reactions that generate orthogonal protein outputs. To identify split ribozymes that require templating for splicing, we used laboratory evolution to evaluate the activities of different split variants of the *Tetrahymena thermophila* ribozyme. The best design delivered a 93-fold dynamic range of splicing with RENDR controlling fluorescent protein production in response to an RNA input. We resolved a thermodynamic model to guide RENDR design, showed how input signals can be transduced into diverse visual, chemical, and regulatory outputs, and used RENDR to detect an antibiotic resistance phenotype in bacteria. This work shows how transcriptional signals can be monitored *in situ* using RNA synthetic biology and converted into different types of biochemical information.

## INTRODUCTION

Cells respond to environmental and intracellular signals by modulating the levels of a wide variety of biomolecules that ultimately determine cell physiology and phenotype. Regulatory mechanisms that operate at the transcriptional level represent a major control point in the process of converting instructions contained in the genome into proteins^1^. RNA thus provides a unique readout of a cell’s identity, physiologic status, and phenotype. The development of genetically-encoded technologies that are able to monitor and transduce individual RNA signals into orthogonal genetic outputs would advance cellular programming such that engineered components can be dynamically coupled with cell state to implement engineered functions^2,3^. Such technologies could, for example, realize genetic programs that are only active in response to species- and cell-specific signatures, link cell states to biochemical processes, and form the foundation for disease-state-sensing therapeutics. Likewise, transducing RNA signals into biomolecular reporters would allow for monitoring of RNA inside of livings cells to uncover quantitative temporal and spatial patterns of gene expression, and allow us to elucidate a deeper understanding of regulatory networks and the processes they control. Compared to monitoring proteins, RNA detection is an attractive option as it can be implemented using easily-designed Watson-Crick base pair interactions, operates on fast timescales^4^, expands the breadth of detection to include non-coding RNA^5^, and it can theoretically access both genotype and phenotype information of host cells. Thus, there is strong motivation to create plug-and-play RNA detection platforms able to transduce intracellular RNA signatures into diverse, orthogonal biomolecular outputs for cell monitoring and genetic programming applications.

In protein engineering, technologies that use protein-protein interactions to control the functionality of an orthogonal protein output have been achieved using topological engineering of split proteins^6,7^. These systems are based on a protein output that is split into two inactive fragments, where interacting protein domains are translationally fused to each fragment. Upon interaction of the fused protein domains, the two fragments of the split-protein output are co-localized and the function of the protein is restored. As a result of the versatility of this approach, a wide range of protein outputs, from enzymes to transcription factors, have been engineered as split proteins^8–14^. Similarly, a variety of protein interaction domains have been used to sense and respond to specific chemical and cellular interactions, yielding an abundance of tools to detect protein-protein interactions^15^, sense small molecules^16^, and implement synthetic gene networks^14^.

When it comes to RNA, programming synthetic RNA to interact with a given RNA signature through Watson-Crick base pairing is relatively straightforward, but transducing this interaction into a biomolecular output in a controlled and orthogonal manner remains challenging. While the potential of using split-RNA reporters and split ribozymes for RNA sensing applications has been previously recognized^17–19^, these technologies typically suffer from limitations that include low-dynamic range (on/off signal), a lack of quantitative design rules to guide the design of new detection interactions, and a lack of modularity for different classes of biomolecular outputs. To address these drawbacks, we have engineered a high-performing, plug-and-play RNA-sensing platform we call Ribozyme ENabled Detection of RNA (RENDR). With RENDR, a cellular RNA input activates a splicing reaction that in turn, produces an mRNA encoding any chosen orthogonal protein output. To achieve this regime, a splicing ribozyme is synthetically split into two non-functional fragments that are each appended with RNA guide sequences which are designed to interact with the RNA input. The split ribozyme is then inserted within a desired gene output. When present, the RNA input co-localizes the two transcribed ribozyme fragments, forming a functional ribozyme complex, and the ribozyme splices together the mRNA of the protein output. To create the RENDR platform, we first characterized nearly all potential split sites across the splicing ribozyme from *Tetrahymena thermophila* by applying for the first time a high-throughput laboratory evolution approach from protein engineering to engineer split RNAs. Using this strategy, we profiled functional split sites across the ribozyme structure and identified novel split designs that allowed for high dynamic range of detection of up to 93-fold. We then investigated the design rules for RNA detection by systematically modifying the interactions between the RNA input and RENDR. By characterizing these variants, we derived a simple thermodynamic model that can guide the design of RENDR variants to transduce new RNA signals. To allow for facile exchange of different genes encoding protein outputs, we developed a modular design strategy that we then harnessed to create RENDR variants that transduce RNA input signals into fluorescent, colorimetric, gaseous, and regulatory outputs. Finally, we used the optimized platform to construct a RENDR device able to sense cellular RNAs that report on the expression of antibiotic resistance genes, demonstrating the utility of this new platform for detecting host cell phenotype.

## RESULTS

### High-throughput identification of ribozyme split sites using *in vitro* transposon mutagenesis coupled with fluorescence-activated cell sorting

To enable convenient measurement of the ribozyme splicing reaction, we first established a fluorescence-based splicing assay in *E. coli*. To do this, a splicing ribozyme from *Tetrahymena thermophila* was inserted within the coding sequence (CDS) of a super folder green fluorescent protein (sfGFP) gene such that translation of the full-length protein is disrupted in the absence of splicing. Upon transcription and splicing, the ribozyme removes itself from the flanking exons and forms a functional sfGFP mRNA, which is then translated to produce a fluorescent protein output (**Fig. 1A**). To decide where to insert the ribozyme within the sfGFP CDS, two design criteria were used: (1) the ribozyme must be inserted immediately downstream of a uracil, which is required for splicing, and (2) the ribozyme should be inserted at a position within sfGFP such that translation of the first exon does not produce a fluorescent signal. Based on these criteria, we constructed a series of plasmids using several ribozyme insertion sites and measured the fluorescence produced by each in *E. coli* (**Supplementary Fig. 2**). From these data, we chose to proceed with the ribozyme inserted after the first nucleotide of amino acid Y66 in the sfGFP CDS (**Fig. 1B and Supplementary Fig. 3**).

**Fig. 1.**
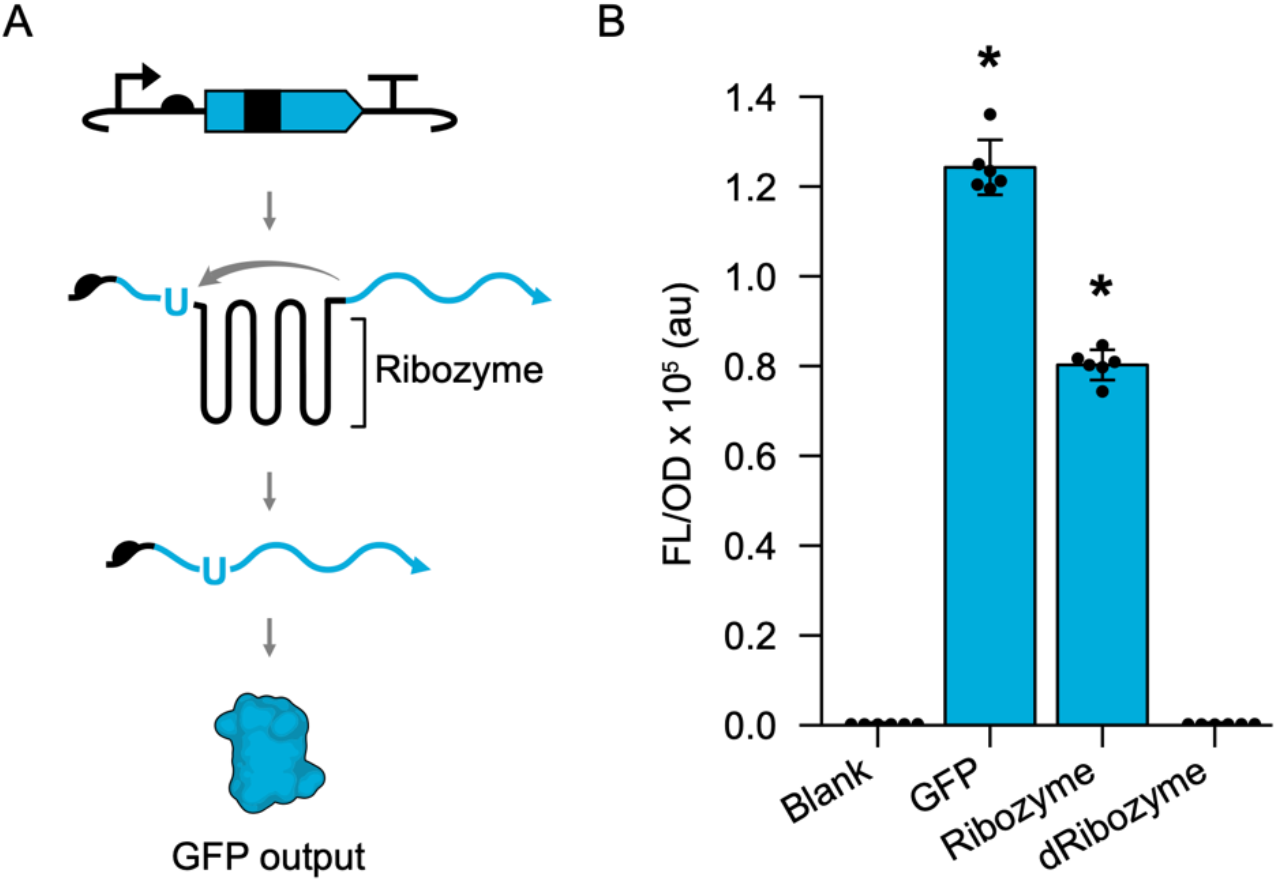
Fluorescence splicing assay enables convenient measurement of ribozyme splicing activity. (**a**) Splicing assay utilizes a plasmid containing an sfGFP coding sequence (blue color) with the *Tetrahymena thermophila* ribozyme DNA sequence (black color) inserted within codon Y66. When this DNA is transcribed into RNA, the ribozyme splices together the sfGFP exons at the 5’ splice site (blue U), located just upstream of the ribozyme, and the spliced product is translated into the native sfGFP (GFP output). (**b**) Fluorescence characterization (measured in units of fluorescence [FL]/optical density [OD] at 600 nm) performed in *E. coli* transformed with an empty plasmid (Blank), a plasmid containing sfGFP (GFP), a plasmid containing sfGFP with an inserted wild-type ribozyme (Ribozyme), and a plasmid containing sfGFP with an inserted catalytically dead G264A mutant ribozyme (dRibozyme). Bars show mean values and error bars represent s.d. of n = 6 biological replicates shown as points. Asterisks indicate p-value < 0.05 relative to blank.

Next, to create a RENDR platform with a high dynamic range of detection, we turned our attention to identifying the optimal ribozyme split sites. An ideal ribozyme split site would destabilize complex formation between the two ribozyme fragments in the absence of RNA input, leading to off states with low signal output. In the presence of the RNA input, this site would have a high level of functional complementation, leading to on states with high signal output. While a few viable split sites within the spicing ribozyme have been identified^20,21^, a systematic analysis of split sites has not been performed. Given the large number of possible split sites (419 variants), we applied a high-throughput, transposon-mediated approach to create a near complete library of split ribozyme variants^6^. While this method has widely been used to optimize split proteins, it has not yet been utilized to engineer split RNAs. In brief, this method uses a Mu transposase to incorporate a synthetic transposon into every position within the ribozyme sequence. Subsequent cloning steps replace the synthetic transposon with complementary RNA interaction sequences, which we call RNA guides, to create split ribozymes (**Fig. 2A and Supplementary Fig. 4**). While our ultimate goal was to create a split-ribozyme system that uses an RNA input to template the splicing reaction, we first elected to use two complimentary RNA guides that directly bring together the split-ribozyme fragments, a reaction we reasoned would be more efficient and produce a stronger output signal. In order to allow for screening in the absence of these interactions, we created a ligand-inducible RNA inhibitor sequence to block the interaction between the RNA guides (**Fig. 2A**). This RNA inhibitor was designed to be the reverse complement of one of the RNA guides and leverages a toehold strategy to favor the disassociation of the functional ribozyme complex. We performed *in vitro* transposon mutagenesis, confirmed the diversity of this library with next-generation sequencing (NGS), co-transformed the library into *E. coli* with a ligand-inducible RNA inhibitor, and performed fluorescence activated cell sorting (FACS) to identify variants with high on and low off states in the absence and presence of the RNA inhibitor, respectively (**Fig. 2A and Supplementary Fig. 5**). Sorted libraries were then sequenced by NGS and the relative enrichment was calculated for each split site (**Fig. 2B and Supplementary Fig. 6**). To provide additional validation of this approach, we performed manual screening of individual colonies after sorting using bulk fluorescence measurements. Individual variants validated in this way were pooled based on their functionality and sequenced to identify functional split sites (**Supplementary Figs. 7-8**). Overlaying these data onto the secondary and tertiary structure of the splicing ribozyme^22,23^, we observed many functional split sites (**Fig. 2C** and **Supplementary Fig. 9**). As has been observed for split proteins, functional split sites appear to be particularly enriched in surface-accessible regions within the ribozyme structure^24^. For example, functional split sites are particularly enriched in the P9 domain, a structure that has been experimentally shown to be solvent-accessible^25^. Similarly, split sites appear to be enriched in regions with low sequence conservation^24^. For example, this family of splicing ribozymes (group I introns) are characterized by a conserved core region composed of the helical domains P4–P6 and P3–P7^26^. In general, we observed these conserved regions to have a lower tolerance for split sites, with the exception of the P6 paired region. Taken together, these results suggest that the splicing ribozyme has a tolerance for split sites in specific regions across its sequence, particularly at sites that are surface accessible with low sequence conservation. Additionally, these results demonstrate the value of applying high-throughput protein engineering methodologies to RNA engineering applications.

**Fig. 2.**
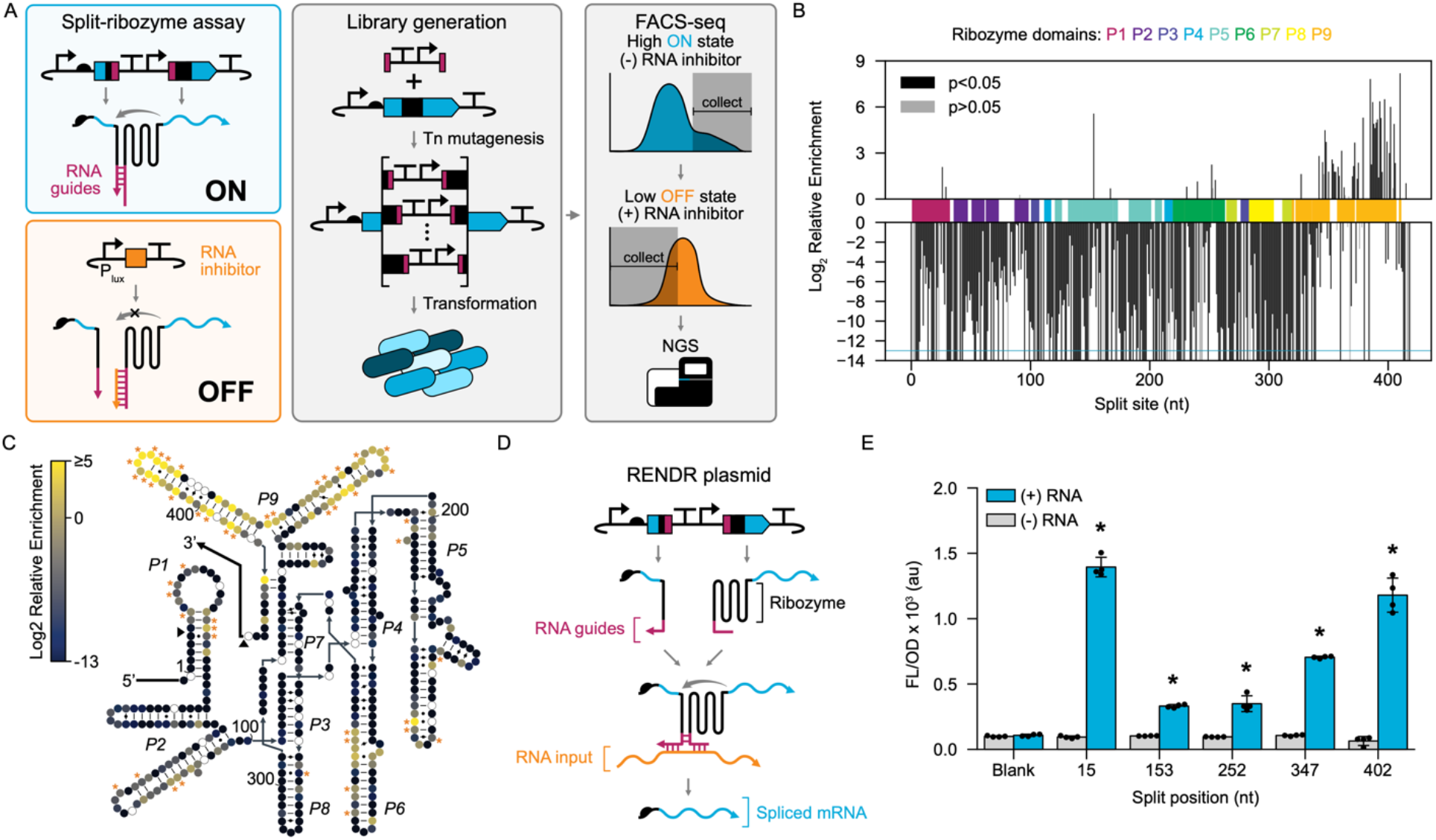
Screening of split ribozymes generates RENDR designs with high dynamic range of detection. (**a**) Schematic of high-throughput screening approach. Left, a split-ribozyme assay is used to determine splicing from each split ribozyme in the absence (on) and presence (off) of additional stabilizing RNA interactions. RNA guides (pink) are used to facilitate RNA interactions between ribozyme fragments, which are blocked with an RNA inhibitor (orange). Center, *in vitro* transposon mutagenesis is used to create a library of split sites. Right, FACS-seq is used to select for split sites with high fluorescence in the on state and low fluorescence in the off state. (**b**) Relative enrichment of ribozyme split sites following FACS-seq (P1-P9 domains and p-values indicated). (**c**) Secondary structure of the ribozyme containing enrichment data for each split site (circle fill color) and split sites identified through colony screening (orange asterisks). Unfilled circles represent split sites absent from the initial library. (**d**) Schematic of RENDR that uses an RNA input to template a split ribozyme, producing a spliced mRNA. (**e**) Characterization of 5 RENDR designs split at identified split sites. Fluorescence characterization (measured in units of fluorescence [FL]/optical density [OD] at 600 nm) was performed in *E. coli* transformed with the RENDR plasmid in the presence and absence of RNA input. Autofluorescence of *E. coli* was determined using cells transformed with empty plasmids (blank). Bars show mean values and error bars represent s.d. of n=4 biological replicates shown as points. Asterisks indicate p-value < 0.05 relative to blank.

### Engineering RENDR devices that transduce cellular RNA signals into detectable outputs

Having identified functional ribozyme split sites, we next aimed to create RENDR devices able to convert cellular RNA signals into orthogonal protein outputs. The basic principle of the RENDR platform is to modify a split ribozyme by appending an RNA guide sequence onto each of the two ribozyme fragments. These RNA guide sequences are designed to base pair with an RNA input - which can be a cellular or synthetic RNA - such that, when present, the RNA input co-localizes the two ribozyme fragments, stabilizes the overall structure, and permits splicing (**Fig. 2D**). To test this, we designed a series of 7 RENDR devices using split sites identified from our data and RNA guides designed to interact with a synthetic RNA input through a total of 163 bp of interaction. Fluorescence characterization revealed that 5 of these RENDR designs were highly functional, leading to higher fluorescence levels in the presence of the RNA input (**Fig. 2E**). We note, that two RENDR designs tested showed a relatively low dynamic range of detection (**Supplementary Fig. 10**). Interestingly, across all designs we observed consistently low off state levels of fluorescence, indistinguishable from autofluorescence of blank cells, in the absence of the RNA input. Likewise, we observed the highest activation for RENDR designs utilizing splits located at the ends of the ribozyme sequence, which we posit could be due to the structural accessibility of these regions. Finally, to confirm that these RENDR designs functioned at the RNA level, as well as to attain a more accurate measure of the dynamic range, we performed RT-qPCR on the high-performing RENDR design, split site 15. We observed a 93-fold increase in the abundance of spliced mRNA in cells expressing both RENDR and RNA input compared to control cells lacking the RNA input (**Supplementary Fig. 11**), a dynamic range comparable to other high-performing synthetic RNA switches^27–29^. Taken together, these results confirm the efficacy of using split ribozymes to create high-performing, RNA signal transduction devices.

### Design rules of RNA input detection

Having explored the design rules governing how to split the catalytic unit of RENDR, we next investigated how the length of the interaction between RNA guides and the RNA input affects the detection reaction. To do this, we constructed five RNA inputs with varying lengths and RENDR devices with corresponding RNA guides. We then used the sfGFP-splicing assay to characterize their performance (**Fig. 3A**). While we observed consistently low off state fluorescence with RENDR devices in the absence of RNA input, there were significant differences in the on state fluorescence values of RENDR devices with different RNA guide lengths. Specifically, we observed an increase in fluorescence as RNA interaction lengths increased up to 164 bp, after which a decrease in fluorescence was observed. To gain a more quantitative insight into this observation, we constructed a simple thermodynamic sequence-function model that uses available RNA free energy and structure prediction tools^30^, and builds upon the success of other RNA mechanistic models^27,31,32^. This model relies on the predicted free energies of several states in the RNA detection reaction (**Fig. 3B**). The initial state (IS) accounts for all RNA species present individually. The seed complex (SC) is formed when the two RNA guides are partially hybridized to the RNA input. The extended duplex (ED) occurs when the RNA guides and the RNA input are fully hybridized. Based on our experimental data and the fast timescales of ribozyme folding and splicing, we hypothesized that SC formation was faster than ED formation and that formation of the SC was sufficient to promote correct complementation and folding of the split ribozyme. With these assumptions, we predicted that the rate of splicing, and hence gene expression, is proportional to the formation of the SC between the RNA input and the two RNA guides (**Supplementary Note 1**).

**Fig. 3.**
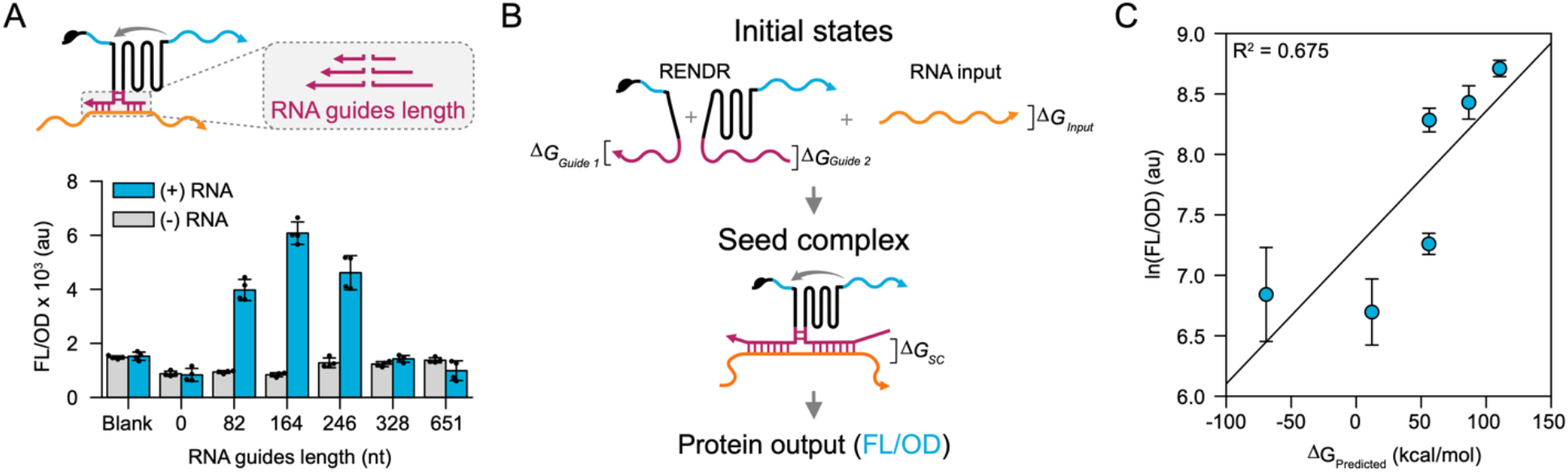
RNA guide length influences RENDR output levels and can be used as a predictive parameter. (**a**) Schematic of RENDR with varying RNA guide lengths and fluorescence characterization of RENDR variants. Fluorescence characterization (measured in units of fluorescence [FL]/optical density [OD] at 600 nm) was performed on *E. coli* transformed with each RENDR design in the presence (blue) and absence (grey) of a plasmid expressing the RNA input. Autofluorescence of *E. coli* was determined using cells transformed with empty plasmids (blank). (**b**) Schematic of a thermodynamic model that considers the free energy of the RNA guides and RNA input in their unbound initial states (IS) and a seed complex (SC) in which they are partially hybridized. Under the hypothesis that the SC is sufficient to stabilize the ribozyme structure and lead to splicing, the natural log of the observed protein output (FL/OD) is proportional to the ΔG_Predicted_, which is the difference in free energies between the IS and the SC. (**c**) Observed correlations between the fluorescence characterization of protein output (natural log FL/OD) and ΔG_Predicted_ of different RNA guide lengths. Coefficient of determination (R^2^) is displayed in the top left corner. Bars in (**a**) and points in (**c**) represent mean values and error bars represent s.d. of n = 4 biological replicates shown as points in (a).

Deriving this model, we predicted that the natural log of the observed gene expression (fluorescence (FL)/optical density at 600 nm (OD)) is linearly related to the difference in free energy between the IS and the SC. For this analysis we only considered folding and interactions of the two RNA guides attached to each ribozyme fragment and the RNA input:

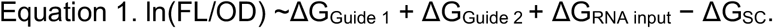

This free energy difference naturally reflects the competing effects of intramolecular base pairs that need to be broken before the formation of intermolecular base pairs that lead to the SC and, ultimately, the active ribozyme state. Comparison of the expression levels predicted by the model to the corresponding experimentally observed fluorescence levels shows strong correlation (R^2^ = 0.68) (**Fig. 3C**). Together, the *in silico* and *in vivo* results suggest that a length of interaction of ~173 bp is optimal for transduction of RNA signals by RENDR and provide a theoretical basis for future design rules and predictive models.

### A modular protein output strategy for RENDR

We next aimed to create a modular gene output design for the RENDR platform that would allow for simple exchange of the output protein being used. To do this, we redesigned the split ribozyme to be located between the RBS and the CDS of the output protein (**Fig. 4A**), resulting in a more modular design that eliminates the need to identify a new location to insert the split ribozyme for every protein-output-coding gene. This design greatly simplifies and generalizes the process of coupling RENDR with different classes of biochemical processes to adapt the platform for different applications. To confirm the functionality of the modular design, we first tested it using a fluorescent sfGFP output (**Fig. 4B**). Following this, we used RENDR to control the biosynthesis of indigo, a chemical commodity that can also serve as a colorimetric output. We combined RENDR with the flavin-containing monooxygenase (FMO) gene from *Methylophaga aminisulfidivorans st. MP*^33^, which catalyzes the conversion of tryptophan-derived indole into indoxyl, which in turn spontaneously forms indigo. Characterization of this design (RENDR-FMO) in *E. coli* revealed tight control of indigo production in the absence of the RNA input, and strong induction of indigo synthesis in the presence of the RNA input (**Fig. 4C and Supplementary Fig. 12**). Next, we aimed to use RENDR to control a gas-producing enzyme, methyl halide transferase (MHT), which produces methyl-bromide (CH3Br) from S-adenosyl methionine (AdoMet) and bromide, and has been used as a reporter system for measuring gene expression in complex and non-transparent matrices like soils^34^. Characterization by GC/MS revealed RENDR-MHT could effectively control the production of CH3Br (**Fig. 4D**). Finally, we aimed to couple RENDR with a CRISPR-Cas system, which is widely used in a variety of gene regulation and editing applications^35,36^. Specifically, we employed a CRISPR interference (CRISPRi) assay, which uses a catalytically dead Cas9 (dCas9) protein to transcriptionally repress a genomically-encoded sfGFP gene, placing translation of the dCas9 under the control of RENDR (RENDR-dCas9). This assay demonstrated strong repression in the presence of the RNA input, suggesting that dCas9 production can be regulated by an RNA signal using RENDR (**Fig. 4E**). We did observe slight repression from RENDR in the absence of the RNA input, implying that RENDR does not fully prevent the expression of dCas9 and that these small levels are sufficient to repress the single sfGFP copy within the *E. coli* genome. Taken together, these results show that RENDR offers a highly modular output design that can transduce cellular RNA input signals into the production of fluorescent, colorimetric, gaseous, and regulatory outputs.

**Fig. 4.**
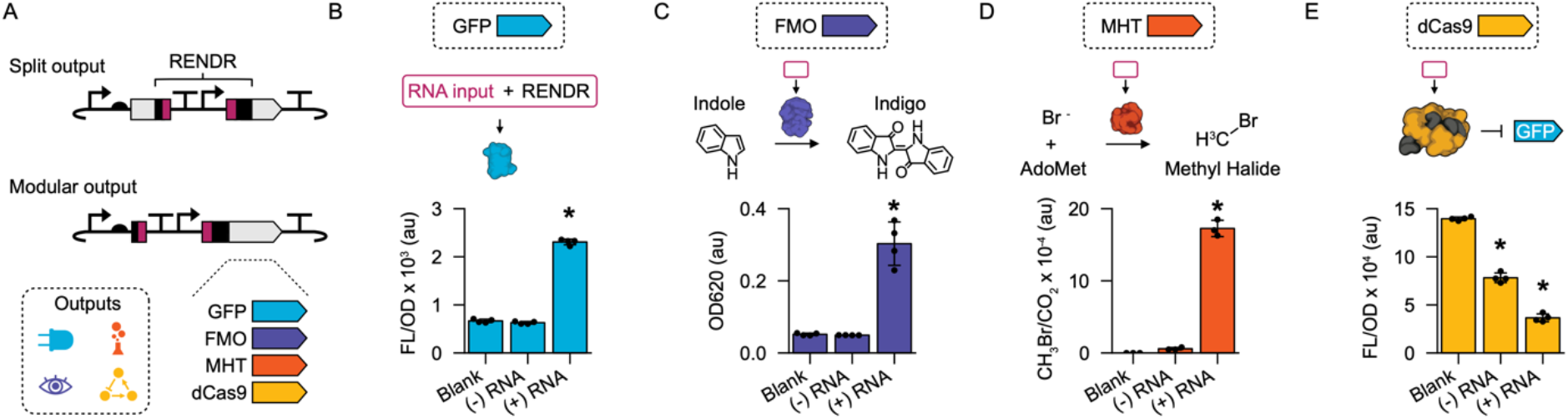
A modular RENDR design allows for interchangeable protein outputs in response to RNA inputs. (**a**) Schematic of a non-modular output design that depends upon gene-specific splits in the CDS (top) and a schematic of a modular output design that uses RENDR to separate the RBS from any CDS output (bottom). The modular design facilitates the simple exchange of different outputs that produce fluorescence, gaseous, colorimetric, and regulatory outputs. (**b**) Fluorescent characterization (measured in units of fluorescence [FL]/optical density [OD] at 600 nm) from RENDR using an sfGFP output. (**c**) Characterization of indigo production (measured in optical density [OD] at 620 nm) from RENDR using a flavin monooxygenase (FMO) output. (**d**) Characterization of methyl-bromine gas production (measured in CH_3_Br/CO_2_) from RENDR using a methyl-halide transferase (MHT) output. (**e**) Characterization of a dCas9 protein output from RENDR measured using an sfGFP-repressing CRISPR interference assay in cells containing a genomically integrated sfGFP expression cassette. For (**b**-**e**), characterization was performed in *E. coli* transformed with each RENDR design in the presence (+) and absence (−) of a plasmid expressing the RNA input. Background signal from *E. coli* was determined using cells transformed with empty plasmids (blank) for each output. In (e), blank uses *E. coli* containing chromosomally integrated sfGFP expression cassette. Bars show mean values and error bars represent s.d. of n = 4 biological replicates shown as points. Asterisks indicate p-value < 0.05 relative to blank.

### Using RENDR to detect antibiotic resistance phenotype

The rise in antibiotic resistant bacteria is a major threat facing public health. One of the major routes that bacteria can acquire antibiotic resistance is through horizontal gene transfer of plasmids encoding antibiotic resistance genes^37^, which can rapidly spread between diverse species across microbial consortia. We posited it would be possible to use RENDR to detect antibiotic resistant phenotypes in bacteria by sensing the mRNAs encoding antibiotic resistance genes (**Fig. 5A**). To test this, we constructed a RENDR device that detects an mRNA encoding for a kanamycin resistance gene (kanR) expressed from a plasmid. This RENDR device (kanR-RENDR) was designed to form 164 bp of interaction with the kanR mRNA and produce an sfGFP output using the modular output design. Transformation of the kanR-RENDR device into cells lacking the KanR-expressing plasmid produced a low fluorescent signal that was comparable to the autofluorescence of blank *E. coli*, while transformation into cells containing the KanR plasmid produced strong fluorescent signals (**Fig. 5B**). Thus, these results show that RENDR can be used to sense antibiotic resistance phenotypes through detection of native cellular mRNAs.

**Fig. 5.**
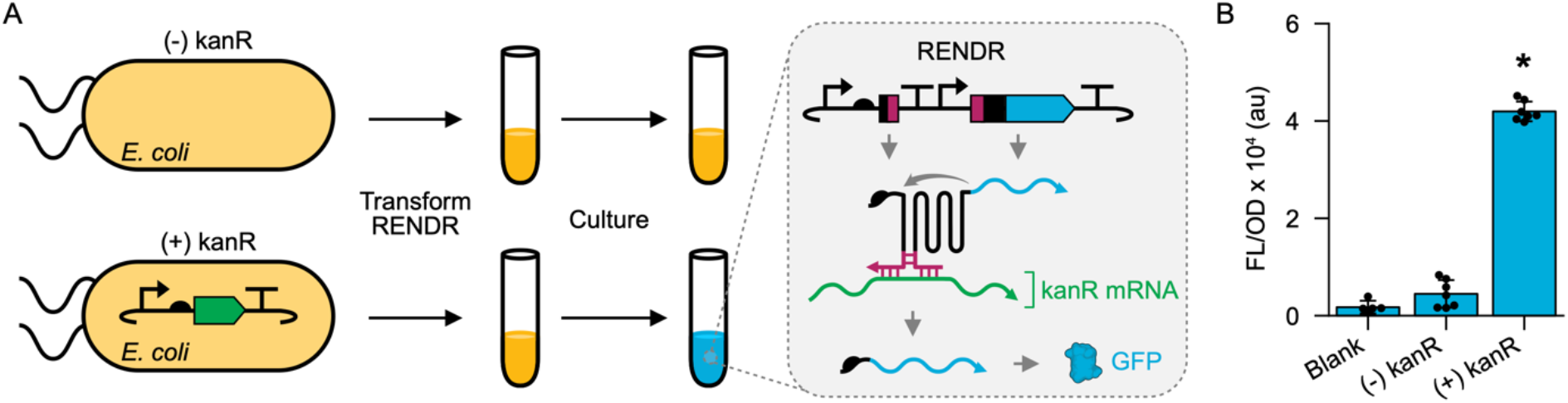
RENDR can identify cells with antibiotic resistance phenotypes. (**a**) Schematic of using RENDR to detect kanamycin resistant cells. RENDR was transformed into E. coli containing (+kanR) or lacking (-kanR) a plasmid expressing a kanamycin resistance gene. When kanR mRNA is present, it is able to activate RENDR, allowing for production of an sfGFP output. (**b**) Fluorescence characterization (measured in units of fluorescence [FL]/optical density [OD] at 600 nm) was performed in *E. coli* transformed with the RENDR plasmid in the presence and absence of a plasmid expressing the kanR mRNA input. Autofluorescence of *E. coli* was determined using cells transformed with empty plasmids (blank). Bars show mean values and error bars represent s.d. of n = 7 biological replicates shown as points. Asterisks indicate p-value < 0.05 relative to blank.

## DISCUSSION

Here we show the development and characterization of RENDR, a ribozyme-based platform that uses individual cellular RNA to template splicing reactions, which in turn produce orthogonal protein outputs. Inspired by protein engineering efforts, we used a laboratory evolution approach to construct a library of split-ribozyme variants and identify high-performing split sites using a high-throughput FACS-seq screen. To allow for easy redesign of RENDR, we determined the optimal RNA guide length and derived a thermodynamic model to further characterize the RENDR splicing reaction. We then demonstrated the modularity of the system by repositioning the split ribozyme upstream of the CDS and replacing sfGFP with other protein-coding sequences, yielding an array of fluorescent, colorimetric, gaseous, and regulatory outputs. Thus, RENDR is a highly modular, designable, and high-performing platform that addresses the limitation of existing split-aptamer and split-ribozyme RNA detection platforms^18^. Finally, we used our optimized system to detect native mRNA that encodes an antibiotic resistance protein, aiding in the identification of antibiotic-resistant cells. This work shows how specific transcripts can be detected *in situ* using an RNA synthetic biology tool and converted into different types of biochemical information.

RENDR delivers a high-performing, plug-and-play RNA detection platform that complements current technologies for transducing RNA signals for cellular programming applications. For example, RNA-responsive CRISPR-Cas single guide RNAs (sgRNAs) have been engineered to form self-inhibiting or inactive structures that can be released upon interaction with a target RNA^38–41^, providing a route for coupling CRISPR-Cas activity to a given RNA target. While powerful, their broad utility is limited because these RNA-responsive sgRNAs can only directly control the activity of the corresponding Cas protein. Our RENDR platform addresses this limitation by allowing for the control of theoretically any protein sequence, which can be easily interchanged using our modular output strategy. Additionally, RENDR could be used in conjunction with RNA-responsive sgRNAs to exert an additional layer of control. For example, a multiple RNA input AND gate can be generated by pairing current RNA-regulated sgRNA technologies with RENDR-activated Cas proteins. This implementation, and other combinations of RNA tools, could be impactful in adding regulatory specificity to these systems and reducing Cas-derived burden on cells.

The toehold switch, another RNA-sensing device, allows access to the RBS, and consequently translation, of an mRNA sequence in response to an RNA target^29^. Like RENDR, toehold switches can control a diversity of protein outputs and offer a high-dynamic range of sensing. However, toehold switch designs require the addition of amino acids to the start of the CDS, which could interfere with protein structure-function and consequently limit the modularity of these systems. RENDR is not bound by this requirement and may be better suited to regulate a wide variety of protein outputs, Additionally, unlike translation-based RNA-sensor technologies, RENDR is not limited to protein outputs and can in principle be coupled to outputs mediated by nucleic acids (e.g., CRISPR RNAs, non-coding small RNA regulators, and fluorescent aptamers). Finally, unlike bacterial transcriptional and translational regulators that are potentially challenging to implement across different cell types^42^, splicing ribozymes are known to function in organisms in all domains of life including bacteria, yeast, and mammalian cells^43,44^. We thus anticipate future versions of RENDR to be utilized in a wide variety of organisms. Taken together, RENDR expands and complements the RNA synthetic biology toolbox for transduction of native RNA for cellular programming.

Within the field of RNA synthetic biology, topological engineering of large RNA represents a currently untapped route to diversify and optimize this biomolecule. To date, most synthetic RNA technologies have focused on the rational and computational design of small, non-coding RNA regulators such as small RNA, riboswitches, small self-cleaving ribozymes, and CRISPR RNA^45^. Here we highlight the potential of engineering large catalytic RNA for synthetic biology applications using laboratory evolution. Specifically, we demonstrate how theoretical and experimental approaches from protein engineering can be easily leveraged to rapidly engineer structurally complex RNA for new capabilities. For example, while *in vitro* transposon mutagenesis coupled with high-throughput screening has been widely applied to create split, fused, and circularly permutated proteins^6^, our work represents the first attempt to systematically apply this method for engineering synthetic RNA functions. While we have demonstrated the power of applying these approaches towards creating RNA-sensing split splicing ribozymes, we expect that this is just the beginning. Circular permutation of RNA, for example, would offer distinct topological modifications, which in protein engineering have been used to yield proteins that are optimized for fusion^46^, have synthetic sensing capabilities^47^, or offer enhanced and expanded functional characteristics compared to wild-type topologies^48,49^. Similarly, topological engineering could be applied to other natural and synthetic ribozymes offering different chemistries or catalytic properties^50^. Thus, in the coming years we envision topological engineering of large catalytic RNA to be used to further expand the structure-function landscape of RNA tools.

Dynamic monitoring of RNA in living cells is a valuable sensing modality. Understanding temporal patterns of transcription is critical to uncovering a deeper understanding of gene regulatory networks and the processes they control^51^. While this can be achieved by labelling endogenous genes with fluorescent markers (e.g., fluorescent proteins or aptamers), RENDR provides an alternative approach to achieve this that circumvents the need for time-consuming and arduous genome engineering. Compared to other RNA templated detection reactions such as fluorescence in situ hybridization (FISH) that depend upon the use of fixed cells, RENDR can be deployed in living cells and allows for dynamic measurements. Beyond direct RNA monitoring, we anticipate RENDR will see broad utility for implementing genetic programs in response to cell phenotype, physiology, and identity. For example, in bacteria, coupling genetic programs to species-specific RNA signatures using RENDR would provide a route to create genetic programs that only function in a desired target host, a goal that is difficult to achieve using traditional regulatory elements (e.g., promoters and RBSs) that often function across related species. Similarly, by coupling genetic programs to specific RNA signatures, we anticipate it will be possible to use RENDR to implement burden- and growth-driven feedback within genetic programs. Such feedback control systems are of critical importance to achieve robustness in engineered functions under fluctuating demands for resources and differing environmental conditions^52^, and to optimize metabolic processes^53^. Finally, we anticipate that RENDR could be an impactful tool for mammalian synthetic biology to, for example, create therapeutic genetic programs that sense and respond to biomarkers of disease^54^. Taken together, we predict that the broad utility of RENDR will advance both basic science and application-driven research in the coming years.

## METHODS

### Plasmid assembly and strains

All plasmids used in this study are listed in **Supplementary Table 1**, with key sequences provided in **Supplementary Tables 2-5 and Supplementary Fig. 1**. *E. coli* strains used are listed in **Supplementary Table 6**. Methods used to clone plasmids included inverse PCR, Gibson Assembly, and Golden Gate assembly. All plasmid constructs were verified using sanger sequencing.

### Fluorescence measurements

Fluorescence measurements were performed in various transformed *E. coli* strains according to **Supplementary Table 6** and as described below. Experiments included quadruplicate measurements of each strain unless otherwise stated. In each experiment, plasmids were transformed into chemically competent *Escherichia coli* str. K-12 substr. MG1655 (*E. coli* MG1655) or TG1 cells, plated on LB-agar plates containing the appropriate antibiotics (LB-agar-Ab), and incubated overnight at 37 °C. Antibiotics were used at the following concentrations: spectinomycin (50 μg/mL), kanamycin (100 μg/mL), carbenicillin (100 μg/mL), and chloramphenicol (34 μg/mL). Following incubation, colonies were used to inoculated 300 μL of LB media containing appropriate antibiotics (LB-Ab) in a deep 96-well plate and incubated at 37 °C at 1000 rpm overnight. Following incubation, 10 μL of each overnight culture was used to inoculate 290 μL of pre-warmed LB-Ab and grown at 37 °C and 1000 rpm for 8 hours (hr). For experiments using N-(β-ketocaproyl)-L-Homoserine lactone (AHL)- and arabinose-inducible plasmids, 4 μL of overnight culture was added to 296 μL of pre-warmed LB-Ab and grown at 37 °C at 1000 rpm for 4 hr. This pre-culture was then diluted 1:50 in fresh, pre-warmed LB-Ab with either 1 μM AHL or 1 % (w/v) arabinose (Alfa Aesar) and grown at 37 °C at 1000 rpm for 6 hr. Bulk fluorescence measurements were performed on 50 μL of experimental culture diluted in 50 μL of phosphate buffered saline (PBS) in a 96-well plate. Optical density (OD) at 600 nm and sfGFP fluorescence (FL) (excitation: 485 nm, emission: 520 nm) was measured using a Tecan M1000 plate reader. For flow cytometry measurements, cultures were diluted 1:100 in PBS and analyzed on a Sony Biotechnology SH800 Cell Sorter. 50,000 events were collected unless otherwise noted.

### Bulk fluorescence data analysis

Each 96-well block included at least two sets of controls; a media blank and *E. coli* transformed with combinations of empty control plasmids pJEC101, pJEC102, or pJEC103, referred to here as blank cells. Blank cells do not express reporter genes, only antibiotic resistance genes, and were used to determine autofluorescence levels. OD and FL values for each colony were first corrected for by subtracting the mean value of the media blank from the respective value of the experimental conditions. The ratio of the corrected FL to the corrected OD (FL/OD) was then calculated for each well.

### Split-ribozyme library

The library of split-ribozyme variants was created using a previously described Mu transposase method^6,55^ (**Supplementary Fig. 4**). In brief, transposon DNA was prepared by digesting 2 μg of plasmid pBW001 (addgene #131529) with HindIII and Bglll at 37 °C for 6 hr, followed by purification. Transposon insertion reactions were then prepared using 50 fmol of purified transposon, 100 fmol of a ribozyme-containing plasmid, 5x MuA buffer, and 0.22 μg of MuA transposase (ThermoFisher Scientific). The reaction was incubated at 30 °C for 16 hr, followed by heat inactivation at 75 °C for 10 minutes and purification. Purified transposon-inserted ribozyme library was then transformed into chemically competent *E. coli* MG1655 cells. A small sample (0.5% of total volume) of the transformation was plated on LB-agar-Ab and the resulting colonies were used to calculate transformation efficiency. The remaining transformation was used to inoculate 50 mL of LB media containing spectinomycin (50 μg/mL) and kanamycin (100 μg/mL), which was grown at 37 °C at 250 rpm overnight, after which library plasmids were isolated using a DNA midi prep kit. Transposon-inserted ribozyme fragments were isolated by digesting 2 μg of transposon-inserted ribozyme plasmids with BbsI-HF at 37 °C for 2 hr, followed by agarose gel purification. The purified transposon-inserted ribozyme variants were then cloned into a plasmid containing a sfGFP gene, which was split after the first nucleotide in amino acid Y66. The transposon was replaced with a linear DNA sequence containing the interacting RNA guide sequences and transformed using the same method as above. Diversity of the cloned library was determined by using PCR to generate amplicons containing the 5’ fragment of the split-ribozyme. After column purification, the PCR product was sent for next generation sequencing (NGS) (Amplicon-EZ, GeneWiz) (**Supplementary Fig. 5**).

### FACS-seq

The split-ribozyme library was co-transformed into *E. coli* MG1655 with plasmid pJEC758, which encodes an AHL-inducible RNA inhibitor. These cells were then subjected to three rounds of fluorescence activated cell sorting (FACS) to enrich for functional split variants. In the first round of sorting, 100 μL of co-transformed library was used to inoculate 5 mL of LB-Ab and grown overnight at 37 °C at 250 rpm. Cells were diluted 1:50 and grown in a pre-culture for 4 hr, then re-diluted 1:50, and grown in culture for 6 hr without AHL. Cells were placed on ice and diluted 1:100 in PBS before being analyzed on a Sony Biotechnology SH800 Cell Sorter. A gate was set to isolate the most fluorescent cells (top 14% of the population) and 673,881 cells were collected in 5 mL LB-Ab. Sorted cells were then incubated at 37 °C at 250 rpm for 1 hr, the volume adjusted to 15 mL using LB-Ab, and grown overnight at 37 °C at 250 rpm. In the second round of sorting, 100 μL of the overnight culture from the first sort were grown in 5 mL LB-Ab using the same protocol used in the first round, with the addition of AHL to the culture at a final concentration of 1 μM for the 6 hr growth period. After growth and dilution in PBS, a gate was set containing the least fluorescent cells (bottom 85% of the population) and 1 million cells were collected in 5 mL LB-Ab. The culture was adjusted to 10 mL with LB-Ab, recovered at 37 °C at 250 rpm for 1 hr, re-adjusted to 15 mL with LB-Ab, and grown overnight at 37 °C at 250 rpm. In the third round of sorting, 100 μL of saturated culture started from cells collected in the second round of sorting were used to inoculate a 5 mL LB-Ab pre-culture and the same protocol was followed that was used in the first round of sorting. A gate was set to isolate the most highly fluorescent cells and 500,000 cells were collected and incubated as in the first round of sorting. A sample of the sorted cells was serially diluted, and plated on LB-agar-Ab overnight at 37 °C. The remaining sorted cells were grown overnight in LB-Ab at 37 °C at 250 rpm, after which glycerol stocks were made and stored at −80 °C.

To perform individual variant screening (**Supplementary Fig. 7**), colonies were picked from serial dilution plates or from plates streaked from glycerol stocks of the sorted library, used to inoculate 300 μL LB-Ab in 96 deep-well plates, and incubated overnight at 37 °C at 1000 rpm. Following overnight incubation, 4 μL of each overnight culture was used to inoculate 196 μL of pre-warmed LB-Ab with or without 0.2 μM AHL and grown for 7 hr at 37 °C at 1000 rpm. After growth, glycerol stocks were made of each culture and duplicate bulk fluorescence measurements performed. We note that some variants were isolated from cultures subjected to a fourth round of sorting following the protocol outlined above, which was used to isolate highly fluorescent cells. From these measurements, each colony was scored based on their on and off states (**Supplementary Fig. 7**). A distance from the x=y line value was calculated for each variant, where Distance = ([FL/OD (-AHL)] - [FL/OD (+AHL)]) / sqrt(2), and variants were split into tertiles based on their distance. Tertiles for each replicate were compared across duplicates and variants in the top two tertiles in both replicates were considered functional. Functionally-validated variants were pooled from glycerol stocks into 45 mL LB-Ab cultures and grown overnight at 37 °C at 250 rpm. Plasmid DNA of the pooled culture was isolated using a Midiprep Kit (Qiagen) and PCR was used to generate amplicons containing the 5’ fragment of the split-ribozyme. After column purification, the PCR product was sent for NGS (Amplicon-EZ, GeneWiz) **(Supplementary Fig. 8B)**.

### Next generation sequencing data analysis

FASTQ files from NGS were processed using a custom Python script described in **Supplementary Note 2**.

### Calculation of free energies for guide length variants

A locally-installed version of the Nucleic Acids Package (NUPACK) version 4.0.0.27 ^56^ was used to calculate free energy parameters for the thermodynamic model (**Supplementary Note 1**).

### FMO output characterization

Plasmids were transformed as described for fluorescence measurements. Colonies (biological quadruplicates) were used to inoculate 300 μL of LB-Ab in a 96 deep-well plate overnight at 37 °C at 1000 rpm. For each overnight culture, 100 μL was transferred to a 5 mL LB-Ab culture containing 5 mM tryptophan and grown to saturation at 37 °C at 250 rpm for 24 hr. Indigo was extracted from cultures using dimethyl sulfoxide (DMSO) ^57^. Briefly, 1.5 mL of each cell culture was centrifuged for 10 minutes at 13,000 g. Pellets were resuspended in 100 μL water, 1 mL of DMSO was added, and samples were vortexed. To quantify the yield, 100 μL of extracted indigo was added to a 96-well plate and the OD at 620 nm was measured for all samples.

### MHT output characterization

Plasmids were transformed as described for fluorescence measurements. Colonies (biological triplicates) were used to inoculate 50 mL tubes with 5 mL LB-Ab and grown overnight at 37 °C at 250 rpm. Cells were washed and normalized to a fixed OD before being transferred to GC-MS vials. Briefly, 1 mL of each overnight culture washed three times by centrifugation at 13,000 g for 1 min followed by resuspension in 1 mL M63 medium (250 μL 1 M MgSO_4_, 2.5 mL 20% glucose, 25 μL 0.5% thiamine, 1.25 mL 10% Bacto casamino acids, 250 μL 1000x spectinomycin, 250 μL 1000x carbenicillin, 6.25 mL 4 M NaBr, 50 mL 5x M63 salts [10 g (NH_4_)_2_SO_4_, 68 g KH_2_PO_4_, 0.00025 g FeSO_4_.7H_2_O, 30 mL 6 M KOH to pH = 7, water up to 1 L], and water up to 250 mL). Washed cells were diluted to OD 600 = 0.05 with M63 medium in a final volume of 1 mL in 2 mL Phenomenex glass vials and subsequently crimped every 2.5 minutes to account for the time it takes the Gas Chromatography/Mass Spectrometry (GC/MS) instrument to process each sample. Crimped glass vials were grown at 37 °C at 250 rpm for 6 hr. A GC/MS containing an Agilent 8890 gas chromatograph fit with an Agilent 7693A autosampler and a 5977B mass spectrometer was used to measure samples. The Agilent MassHunter Workstation Quantitative Analysis software was used to quantify resulting methyl-bromide and carbon dioxide (CO_2_) production. To account for cell density, the methyl-bromide production was normalized to CO_2_ concentration for each sample.

### Statistical analysis

The sample average and standard deviation (s.d.) were calculated from the replicates of each sample in all experiments. Significance in all experiments was assessed using a homoscedastic two-sample t-Test with an alpha of 0.05. In all cases, resulting p-values of less than 0.05 for experimental sample measurements compared to blank cell measurements were considered significantly different.

### Total RNA extraction for reverse transcription quantitative PCR (RT-qPCR)

For RNA extraction and quantitative PCR measurements, *E. coli* strain MG1655 was used and performed in biological triplicate. Plasmids were transformed and colonies were used to inoculate 5 mL of LB-Ab and incubated at 37 °C at 250 rpm overnight. Following incubation, 100 μL of each overnight culture was used to inoculate 5 mL of pre-warmed LB-Ab and grown for 6 hr. After growth, 500 μL of each culture was centrifuged at 13,000 g for 1 min and the supernatant removed. Pellets were resuspended in 750 μL of Trizol reagent (Thermo Fisher Scientific) and incubated at room temperature for 5 min, then 150 μL of chloroform was added, mixed by vortexing for 15 sec, and incubated at room temperature for 3 min. After incubation, samples were centrifuged for 15 min at 12,000 g at 4 °C and the top aqueous layer isolated and 1 μL of RNA-grade glycogen (Thermo Fisher Scientific, 20 mg/mL) and 375 μL of isopropanol were added. The solution was then incubated at room temperature for 10 min and centrifuged at 4 °C at 13,000 g for 15 min. After centrifugation, the isopropanol was carefully removed, pellets were washed in 600 μL of chilled 70% ethanol, and samples were centrifuged at 4 °C at 13,000 g for 2 min. Following removal of the ethanol, the pellets were centrifuged again at 4 °C at 13,000 g for 2 min to ensure the complete removal of the ethanol. Pellets were air dried for 5 min and resuspended in 18 μL of RNase-free water.

### DNase treatment of total RNA for RT-qPCR

The concentration of all samples was measured with a Qubit fluorometer (Thermo Fisher Scientific). For each sample, 1.5 μg of total RNA (for a final concentration of 30 ng/μL) was digested by Turbo DNase (Thermo Fisher Scientific) according to the manufacturer’s instructions. After digestion, 150 μL of RNase-free water and 200 μL of phenol-chloroform were added to each tube. The samples were then mixed by vortexing for 10 sec, incubated at room temperature for 3 min, and centrifuged at 13,000 g at 4 °C for 10 min. After centrifugation, 190 μL of the top aqueous layer was removed and added to 190 μL of chloroform. Samples were vortexed for 10 sec, incubated for 3 min at room temperature, and centrifuged at 13,000 g at 4 °C for 10 min. After centrifugation, 170 μL of the top aqueous layer was removed and added to 170 μL of chloroform. Samples were vortexed for 10 sec, incubated for 3 min at room temperature, and centrifuged at 13,000 g at 4 °C for 10 min. After centrifugation, 120 μL of the top aqueous layer was removed and mixed with 1 μL of RNA-grade glycogen, 360 μL of chilled 100% ethanol, and 12 μL of 3 M sodium acetate. Samples were mixed by pipetting and incubated at −80 °C for at least 1 hr. After incubation, the samples were centrifuged for 30 min at 13,000 g at 4 °C. After removing the supernatant, the resulting RNA pellets were washed with 600 μL of chilled 70% ethanol and centrifuged at 4 °C at 13,000 g for 10 min. After centrifugation the ethanol was removed, and the pellets were centrifuged at 4 °C at 13,000 g for 2 min. Ethanol wash was repeated a second time to ensure the complete removal of any remaining ethanol. Pellets were air dried for 5 min and resuspended in 12 μL of RNase-free water.

### Reverse transcription and qPCR measurements

The concentration of all samples was measured with a Qubit fluorometer. To anneal primers for cDNA synthesis reactions, 75 ng of total RNA, 0.5 μL of 2 μM reverse transcription primer, 1 μL of 10 mM dNTPs (New England BioLabs), and RNase-free water (up to 6.5 μl) were combined. This mixture was then incubated at 65 °C for 5 minutes and cooled on ice for 5 min. For reverse transcription, 0.25 μL of Superscript III reverse transcriptase (Thermo Fisher Scientific), 1 μL of 100 mM Dithiothreitol (DTT), 1x first-strand buffer (Thermo Fisher Scientific), 0.5 μL RNaseOUT (Thermo Fisher Scientific), and RNase-free water up to 3.5 μL were then added to the annealed primer reaction, and the total mixture was incubated at 55 °C for 1 hr, 75 °C for 15 min, and then stored at −20 °C. The resulting cDNA samples were subsequently analyzed via qPCR. Each biological sample for each condition was analyzed in three technical replicates in a 96-well microplate covered with an optically clear seal. Total RNA samples were also analyzed as non-reverse transcription controls (NRT) to confirm that no DNA could be detected under the conditions used. Additionally, a non-template control was run to confirm that no non-specific amplification was present. For each technical replicate, a reaction was prepared by mixing 5 μL of Maxima SYBR green qPCR master mix (Thermo Fisher Scientific), 2 μL of cDNA and 0.5 μL of 2 μM forward and reverse sfGFP qPCR primers, and RNase-free water up to 10 μL. For the reaction, the following program was used: 50 °C for 2 min, 95 °C for 10 min, followed by 30 cycles of 95 °C for 15 s and 60 °C for 1 min. Melting curve analysis was run after all measurements to ensure that only a single product was amplified. For quantification, a standard curve covering four ten-fold serial dilutions of an sfGFP standard was analyzed in parallel. The standard curve was used to determine the relative abundance of sfGFP cDNA in all tested samples and to infer the efficiency of the qPCR primer set (85%). Results were analyzed using CFX Maestro software (Biorad). Briefly, the standard curve was used to determine the starting quantity (SQ) of the sfGFP cDNA in each reaction. The SQ values of technical replicates for each sample were then averaged to give a single value for each biological replicate. The mean and s.d of biological replicates were then calculated for each sample type.

## Acknowledgements

The authors acknowledge Dr. Joff Silberg and the Chappell Lab members for helpful discussion. We would like to especially thank Li Chieh Lu and Malyn Selinidis for their help with the MHT experimental design and analysis. This material is based on work supported by the National Science Foundation Graduate Research Fellowship Program (grant no. 1842494 to L.G), the National Science Foundation (grant no. 2128370 to J.C), and the Welch Foundation (grant no. C-1982-20190330 to JC). J.C. is an Alfred P. Sloan Research Fellow.

## Author contributions

L.G. and J.C. conceived of the project. All authors helped design the experiments, collect data, and write the manuscript.

## Materials & correspondence

Correspondence should be directed to Dr. James Chappell

