## Supplementary information for "A split ribozyme that links detection of a native RNA to orthogonal protein outputs"

### Table of contents

| Tables | Title | Page |
|---|---|---|
| 1 | Plasmids used in this study | 2 - 5 |
| 2 | Example DNA plasmid sequence | 5 - 6 |
| 3 | RNA inputs used in this study | 6 - 8 |
| 4 | Guide sequences used in this study | 8 - 9 |
| 5 | Sequence parts used in this study | 9 - 15 |
| 6 | Strains used in this study | 15 |
| **Figures** | | |
| 1 | Schematic of representative DNA plasmids used in this study | 16 |
| 2 | Identifying functional ribozyme insertion sites in sfGFP | 17 |
| 3 | Single cell fluorescence characterization of fluorescence-based splicing assay | 18 |
| 4 | Schematic of split-ribozyme library generation using transposon mutagenesis | 19 |
| 5 | Diversity of the split-ribozyme library | 20 |
| 6 | FACS of the split-ribozyme library results in enrichment at specific split sites | 21 |
| 7 | Identification of functional split ribozymes | 22 |
| 8 | Comparison of split variants identified through FACS-seq and individual colony screening | 23 |
| 9 | Three-dimensional structure shows functional ribozyme split sites in surface accessible regions | 24 |
| 10 | Two enriched split sites resulted in RENDR variants with low-fold activation | 25 |
| 11 | Reverse transcription quantitative PCR (RT-qPCR) of RENDR spliced output in the presence and absence of RNA input | 26 |
| 12 | RENDR variant using a flavin-containing monooxygenase (FMO) output produces visually discernable indigo levels upon detection of RNA | 27 |
| **Notes** | | |
| 1 | Encapsulating RENDR design principles in a thermodynamic model | 28 - 29 |
| 2 | NGS data analysis pipeline | 29 - 30 |

**Supplementary Table 1. All plasmids used in this study.** CmR: chloramphenicol resistance gene, AmpR: ampicillin resistance gene, SpecR: spectinomycin resistance gene, p15A: origin of replication, ColE1: origin of replication, CDF: CloDF origin of replication, P: promoter, T: terminator, HP14: stability hairpin, RBS: ribosome binding site, P1_loop: P1 domain in *T. thermophila* ribozyme, IGS: internal guide sequence, sfGFP(X-YntZ) = sfGFP gene from amino acid X to amino acid Y at nucleotide position Z. Figure column refers to the figure (F) or supplementary figure (SF) the plasmid was used in. If All, the plasmid was used in every figure.

| Plasmid | Name | Plasmid Architecture | Figure |
|---|---|---|---|
| pBW001 | Transposon | AmpR - ColE1 - Transposon (containing KanR) | SI4 |
| pJEC101 | Empty | CmR - p15A - TrrnB | All |
| pJEC102 | Empty | AmpR - ColE1 - PJ23119 - TrrnB | All |
| pJEC103 | Empty | SpecR - CDF - PJ23119 - TrrnB | All |
| pJEC752 | GFP | SpecR - CDF - PJ23119 - HP14 - RBS_01 - sfGFP(1-239) - TrrnB | F1 |
| pJEC753 | Ribozyme | SpecR - CDF - PJ23119 - HP14 - RBS_01 - sfGFP(1-66nt1) - P1_loop_WT - IGS_01 - Ribozyme - sfGFP(66nt2-239) - TrrnB | F1 |
| pJEC754 | Catalytically dead ribozyme | SpecR - CDF - PJ23119 - HP14 - RBS_01 - sfGFP(1-66nt1) - P1_loop_WT - IGS_01 - G264A dRibozyme - sfGFP(66nt2-239) - TrrnB | F1 |
| pJEC758 | AHL-inducible RNA inhibitor | AmpR - ColE1 - PTet - RBS_02 - LuxR - PLuxR - RNA inhibitor - T500 | F2 |
| pJEC759 | RENDR split 15 nt into ribozyme (guide/input pair 2) | SpecR - CDF - PJ23119 - HP14 - RBS_01 - P1_loop_01 - Stem_01 - Guide_03 - T500 - PJ23119 - Guide_04 - Stem_02 - P1_loop_03 - IGS - Ribozyme - sfGFP(66nt2-239) - T500 | F2,3 |
| pJEC760 | RENDR split 26 nt into ribozyme | SpecR - CDF - TL3S2P55 - mCherry2 - RBS_02 - PJ23115 - PJ23119 - HP14 - RBS_01 - sfGFP(1-66nt1) - P1_loop_WT - Stem_01 - Guide_03 - T500 - PJ23119 - Guide_04 - Stem_02 - IGS_01 - Ribozyme - sfGFP(66nt2-239) - T500 | F2 |
| pJEC761 | RENDR split 153 nt into ribozyme | SpecR - CDF - TL3S2P55 - mCherry2 - RBS_02 - PJ23115 - PJ23119 - HP14 - RBS_01 - sfGFP(1-66nt1) - P1_loop_WT - IGS_01 - Ribozyme(1-121) - Stem_01 - Guide_03 - T500 - PJ23119 - Guide_04 - Stem_02 - Ribozyme(122-388) - sfGFP(66nt2-239) - T500 | F2 |
| pJEC762 | RENDR split 252 nt into ribozyme | SpecR - CDF - TL3S2P55 - mCherry2 - RBS_02 - PJ23115 - PJ23119 - HP14 - RBS_01 - sfGFP(1-66nt1) - P1_loop_WT - IGS_01 - Ribozyme(1-220) - Stem_01 - Guide_03 - T500 - PJ23119 - Guide_04 - Stem_02 - Ribozyme(221-388) - sfGFP(66nt2-239) - T500 | F2 |
| pJEC763 | RENDR split 327 nt | SpecR - CDF - TL3S2P55 - mCherry2 - RBS_02 - PJ23115 - PJ23119 - HP14 - RBS_01 - sfGFP(1-66nt1) - P1_loop_WT - IGS_01 - Ribozyme(1-295) - Stem_01 - | F2 |

| | | | |
|---|---|---|---|
| | into ribozyme | Guide_03 - T500 - PJ23119 - Guide_04 - Stem_02 - Ribozyme(296-388) - sfGFP(66nt2-239) - T500 | |
| pJEC764 | RENDR split 347 nt into ribozyme | SpecR - CDF - TL3S2P55 - mCherry2 - RBS_02 - PJ23115 - PJ23119 - HP14 - RBS_01 - sfGFP(1-66nt1) - P1_loop_WT - IGS_01 - Ribozyme(1-315) - Stem_01 - Guide_03 - T500 - PJ23119 - Guide_04 - Stem_02 - Ribozyme(316-388) - sfGFP(66nt2-239) - T500 | F2 |
| pJEC765 | RENDR split 369 nt into ribozyme | SpecR - CDF - TL3S2P55 - mCherry2 - RBS_02 - PJ23115 - PJ23119 - HP14 - RBS_01 - sfGFP(1-66nt1) - P1_loop_WT - IGS_01 - Ribozyme(1-337) - Stem_01 - Guide_03 - T500 - PJ23119 - Guide_04 - Stem_02 - Ribozyme(338-388) - sfGFP(66nt2-239) - T500 | F2 |
| pJEC766 | RENDR split 402 nt into ribozyme | SpecR - CDF - TL3S2P55 - mCherry2 - RBS_02 - PJ23115 - PJ23119 - HP14 - RBS_01 - sfGFP(1-66nt1) - P1_loop_WT - IGS_01 - Ribozyme(1-370) - Stem_01 - Guide_03 - T500 - PJ23119 - Guide_04 - Stem_02 - Ribozyme(371-388) - sfGFP(66nt2-239) - T500 | F2 |
| pJEC767 | Constitutive RNA input | AmpR - ColE1 - PJ23119 - Input_01 - T500 | F2 |
| pJEC768 | RENDR (non-interacting guide/input pair) | SpecR - CDF - PJ23119 - HP14 - RBS_01 - sfGFP(1-66nt1) - P1_loop_01 - Guide_05 - T500 - PJ23119 - Guide_06 - P1_loop_03 - IGS_01 - Ribozyme - sfGFP(66nt2-239) - T500 | F3 |
| pJEC769 | RENDR (guide/input pair 1) | SpecR - CDF - PJ23119 - HP14 - RBS_01 - sfGFP(1-66nt1) - P1_loop_01 - Guide_07 - T500 - PJ23119 - Guide_08 - P1_loop_03 - IGS_01 - Ribozyme - sfGFP(66nt2-239) - T500 | F3 |
| pJEC770 | RENDR (guide/input pair 3) | SpecR - CDF - PJ23119 - HP14 - RBS_01 - sfGFP(1-66nt1) - P1_loop_01 - Guide_09 - T500 - PJ23119 - Guide_10 - P1_loop_03 - IGS_01 - Ribozyme - sfGFP(66nt2-239) - T500 | F3 |
| pJEC771 | RENDR (guide/input pair 4) | SpecR - CDF - PJ23119 - HP14 - RBS_01 - sfGFP(1-66nt1) - P1_loop_01 - Guide_11 - T500 - PJ23119 - Guide_12 - P1_loop_03 - IGS_01 - Ribozyme - sfGFP(66nt2-239) - T500 | F3 |
| pJEC772 | RENDR (guide/input pair 5) | SpecR - CDF - PJ23119 - HP14 - RBS_01 - sfGFP(1-66nt1) - P1_loop_01 - Guide_13 - T500 - PJ23119 - Guide_14 - P1_loop_03 - IGS_01 - Ribozyme - sfGFP(66nt2-239) - T500 | F3 |
| pJEC773 | RNA input (non-interacting guide/input pair) | AmpR - ColE1 - TTonB - araC - PBad - Input_02 - T500 | F3 |
| pJEC774 | RNA input (guide/input pair 1) | AmpR - ColE1 - TTonB - araC - PBad - Input_03 - T500 | F3 |

| | | | |
|---|---|---|---|
| pJEC775 | RNA input (guide/input pair 2) | AmpR - ColE1 - TTonB - araC - PBad - Input_01 - T500 | F3 |
| pJEC776 | RNA input (guide/input pair 3) | AmpR - ColE1 - TTonB - araC - PBad - Input_04 - T500 | F3 |
| pJEC777 | RNA input (guide/input pair 4) | AmpR - ColE1 - TTonB - araC - PBad - Input_05 - T500 | F3 |
| pJEC778 | RNA input (guide/input pair 5) | AmpR - ColE1 - TTonB - araC - PBad - Input_06 - T500 | F3 |
| pJEC779 | Modular RENDR-GFP | SpecR - CDF - PJ23119 - HP14 - RBS_01 - P1_helix_01- P1_loop_01 - Stem_01 - Guide_15 - T500 - PJ23119 - Guide_16 - Stem_02 - P1_loop_03 - IGS_01 - Ribozyme - sfGFP - TrrnB | F4 |
| pJEC780 | Constitutive synthetic RNA input | AmpR - ColE1 - PJ23119 - Input_07 - T500 | F4 |
| pJEC781 | Modular RENDR-FMO | SpecR - CDF - PJ23119 - HP14 - RBS_01 - P1_helix_01- P1_loop_01 - Stem_01 - Guide_15 - T500 - PJ23119 - Guide_16 - Stem_02 - P1_loop_03 - IGS_01 - Ribozyme - FMO - TrrnB | F4 |
| pJEC782 | Modular RENDR-MHT | SpecR - CDF - PJ23119 - HP14 - RBS_01 - P1_helix_01 - P1_loop_01 - Stem_01 - Guide_15 - T500 - PJ23119 - Guide_16 - Stem_02 - P1_loop_03 - IGS_01 - Ribozyme - MHT - TrrnB | F4 |
| pJEC783 | Modular RENDR-dCas9 | SpecR - CDF - PJ23119 - HP14 - RBS_01 - P1_helix_01 - P1_loop_01 - Stem_01 - Guide_15 - T500 - PJ23119 - Guide_16 - Stem_02 - P1_loop_03 - IGS_01 - Ribozyme - dCas9 - TrrnB | F4 |
| pJEC784 | sgRNA | CmR - p15A - PJ23119 - sgRNA_01 - TrrnB | F4 |
| pJEC785 | KanR sensing RENDR-GFP | SpecR - CDF - PJ23119 - HP14 - RBS_01 - P1_helix_01 - P1_loop_01 - Stem_01 - Guide_17 - T500 - PJ23119 - Guide_18 - Stem_02 - P1_loop_03 - IGS_01 - Ribozyme - sfGFP - TrrnB | F5 |
| pJEC786 | KanR RNA input | AmpR - ColE1 - PJ23119 - RBS_01 - Input_08 - T500 | F5 |
| pJEC787 | Ribozyme split at E172 | SpecR - CDF - PJ23119 - HP14 - RBS_01 - sfGFP(1-172nt3) - P1_helix_02 - P1_loop_WT - IGS_02 - Ribozyme - sfGFP(173nt1-239) - TrrnB | SF2 |
| pJEC788 | Catalytically dead ribozyme split at E172 | SpecR - CDF - PJ23119 - HP14 - RBS_01 - sfGFP(1-172nt3) - P1_helix_02 - P1_loop_WT - IGS_02 - dRibozyme - sfGFP(173nt1-239) - TrrnB | SF2 |
| pJEC789 | Ribozyme split at D21 | SpecR - CDF - PJ23119 - HP14 - RBS_01 - sfGFP(1-21nt3) - P1_loop_WT - IGS_03 - Ribozyme - sfGFP(22nt1-239) - TrrnB | SF2 |

| | | | |
|---|---|---|---|
| pJEC790 | Catalytically dead ribozyme split at D21 | SpecR - CDF - PJ23119 - HP14 - RBS_01 - sfGFP(1-21nt3) - P1_loop_WT - IGS_03 - dRibozyme - sfGFP(22nt1-239) - TrrnB | SF2 |
| pJEC791 | Ribozyme FMO output | SpecR - CDF - PJ23119 - HP14 - RBS_01 - P1_helix_01- P1_loop_WT - IGS_01 - Ribozyme - FMO - TrrnB | SF11 |

**Supplementary Table 2.** Example DNA plasmid sequence.

| Name | DNA sequence |
|---|---|
| Example of cis-acting ribozyme inserted within sfGFP, pJEC753 [SpecR - CDF - PJ23119 - HP14 - RBS_01 - sfGFP(1-66nt1) - P1_loop_WT - IGS_01 - Ribozyme - sfGFP(66nt2-239) - TrrnB] | TTATTTGCCGACTACCTTGGTGATCTCGCCTTTCACGTAGTGGACAAATTC<br>TTCCAACTGATCTGCGCGCGAGGCCAAGCGATCTTCTTCTTGTCCAAGATA<br>AGCCTGTCTAGCTTCAAGTATGACGGGCTGATACTGGGCCGGCAGGCGCT<br>CCATTGCCCAGTCGGCAGCGACATCCTTCGGCGCGATTTTGCCGGTTACT<br>GCGCTGTACCAAATGCGGGACAACGTAAGCACTACATTTCGCTCATCGCC<br>AGCCCAGTCGGGCGGCGAGTTCCATAGCGTTAAGGTTTCATTTAGCGCCT<br>CAAATAGATCCTGTTCAGGAACCGGATCAAAGAGTTCCTCCGCCGCTGGA<br>CCTACCAAGGCAACGCTATGTTCTCTTGCTTTTGTCAGCAAGATAGCCAGA<br>TCAATGTCGATCGTGGCTGGCTCGAAGATACCTGCAAGAATGTCATTGCG<br>CTGCCATTCTCCAAATTGCAGTTCGCGCTTAGCTGGATAACGCCACGGAAT<br>GATGTCGTCGTGCACAACAATGGTGACTTCTACAGCGCGGAGAATCTCGC<br>TCTCTCCAGGGGAAGCCGAAGTTTCCAAAAGGTCGTTGATCAAAGCTCGC<br>CGCGTTGTTTCATCAAGCCTTACGGTCACCGTAACCAGCAAATCAATATCA<br>CTGTGTGGCTTCAGGCCGCCATCCACTGCGGAGCCGTACAAATGTACGGC<br>CAGCAACGTCGGTTCGAGATGGCGCTCGATGACGCCAACTACCTCTGATA<br>GTTGAGTCGATACTTCGGCGATCACCGCTTCCCTCATACTCTTCCTTTTTC<br>AATATTATTGAAGCATTTATCAGGGTTATTGTCTCATGAGCGGATACATATT<br>TGAATGTATTTAGAAAAATAAACAAATAGCTAGCTCACTCGGTCGCTACGC<br>TCCGGGCGTGAGACTGCGGCGGGCGCTGCGGACACATACAAAGTTACCC<br>ACAGATTCCGTGGATAAGCAGGGGACTAACATGTGAGGCAAAACAGCAGG<br>GCCGCGCCGGTGGCGTTTTTCCATAGGCTCCGCCCTCCTGCCAGAGTTCA<br>CATAAACAGACGCTTTTCCGGTGCATCTGTGGGAGCCGTGAGGCTCAACC<br>ATGAATCTGACAGTACGGGCGAAACCCGACAGGACTTAAAGATCCCCACC<br>GTTTCCGGCGGGTCGCTCCCTCTTGCGCTCTCCTGTTCCGACCCTGCCGT<br>TTACCGGATACCTGTTCCGCCTTTCTCCCTTACGGGAAGTGTGGCGCTTTC<br>TCATAGCTCACACACTGGTATCTCGGCTCGGTGTAGGTCGTTCGCTCCAA<br>GCTGGGCTGTAAGCAAGAACTCCCCGTTCAGCCCGACTGCTGCGCCTTAT<br>CCGGTAACTGTTCACTTGAGTCCAACCCGGAAAAGCACGGTAAAACGCCA<br>CTGGCAGCAGCCATTGGTAACTGGGAGTTCGCAGAGGATTTGTTTAGCTA<br>AACACGCGGTTGCTCTTGAAGTGTGCGCCAAAGTCCGGCTACACTGGAAG<br>GACAGATTTGGTTGCTGTGCTCTGCGAAAGCCAGTTACCACGGTTAAGCA<br>GTTCCCCAACTGACTTAACCTTCGATCAAACCACCTCCCCAGGTGGTTTTT<br>TCGTTTACAGGGCAAAAGATTACGCGCAGAAAAAAAGGATCTCAAGAAGAT<br>CCTTTGATCTTTTCTACTGAACCGCTCTAGATTTCAGTGCAATTTATCTCTT<br>CAAATGTAGCACCTGAAGTCAGCCCCATACGATATAAGTTGTAATTCTCAT<br>GTTAGTCATGCCCCGCGCCCACCGGAAGGAGCTGACTGGGTTGAAGGCT<br>CTCAAGGGCATCGGTCGAGATCCCGGTGCCTAATGAGTGAGCTAACTTAC<br>ACCCGCAGTTCCAGTACGGTTCCATTAATTGCGTTGCGCTGCTTCCGGCT |

| | | |
|---|---|---|
| | | GAATTCTAAAGATCTTTGACAGCTAGCTCAGTCCTAGGTATAATACTAGTAC<br>GTCGACTCTCGAGTGAGATTGTTGACGGTACCGTATTTTGGATCTAGGAG<br>GAAGGATCTATGAGCAAAGGAGAAGAACTTTTCACTGGAGTTGTCCCAATT<br>CTTGTTGAATTAGATGGTGATGTTAATGGGCACAAATTTTCTGTCCGTGGA<br>GAGGGTGAAGGTGATGCTACAAACGGAAAACTCACCCTTAAATTTATTTGC<br>ACTACTGGAAAACTACCTGTTCCGTGGCCAACACTTGTCACTACTCTGACC<br>TAAATAGCAATATTTACCTTTGGGTCAAAAAGTTATCAGGCATGCACCTGGT<br>AGCTAGTCTTTAAACCAATAGATTGCATCGGTTTAAAAGGCAAGACCGTCA<br>AATTGCGGGAAAGGGGTCAACAGCCGTTCAGTACCAAGTCTCAGGGGAAA<br>CTTTGAGATGGCCTTGCAAAGGGTATGGTAATAAGCTGACGGACATGGTC<br>CTAACCACGCAGCCAAGTCCTAAGTCAACAGATCTTCTGTTGATATGGATG<br>CAGTTCACAGACTAAATGTCGGTCGGGGAAGATGTATTCTTCTCATAAGAT<br>ATAGTCGGACCTCTCCTTAATGGGAGCTAGCGGATGAAGTGATGCAACAC<br>TGGAGCCGCTGGGAACTAATTTGTATGCGAAAGTATATTGATTAGTTTTGG<br>AGTACTCGATGGTGTTCAATGCTTTTCCCGTTATCCGGATCACATGAAACG<br>GCATGACTTTTTCAAGAGTGCCATGCCCGAAGGTTATGTACAGGAACGCA<br>CTATATCTTTCAAAGATGACGGGACCTACAAGACGCGTGCTGAAGTCAAGT<br>TTGAAGGTGATACCCTTGTTAATCGTATCGAGTTAAAGGGTATTGATTTTAA<br>AGAAGATGGAAACATTCTTGGACACAAACTCGAGTACAACTTTAACTCACA<br>CAATGTATACATCACGGCAGACAAACAAAAGAATGGAATCAAAGCTAACTT<br>CAAAATTCGCCACAACGTTGAAGATGGTTCCGTTCAACTAGCAGACCATTA<br>TCAACAAAATACTCCAATTGGCGATGGCCCTGTCCTTTTACCAGACAACCA<br>TTACCTGTCGACACAATCTGTCCTTTCGAAAGATCCCAACGAAAAGCGTGA<br>CCACATGGTCCTTCTTGAGTTTGTAACTGCTGCTGGGATTACACATGGCAT<br>GGATGAGCTCTACAAATAAGGATCTGAAGCTTGGGCCCGAACAAAAACTC<br>ATCTCAGAAGAGGATCTGAATAGCGCCGTCGACCATCATCATCATCATCAT<br>TGAGTTTAAACGGTCTCCAGCTTGGCTGTTTTGGCGGATGAGAGAAGATTT<br>TCAGCCTGATACAGATTAAATCAGAACGCAGAAGCGGTCTGATAAAACAGA<br>ATTTGCCTGGCGGCAGTAGCGCGGTGGTCCCACCTGACCCCATGCCGAA<br>CTCAGAAGTGAAACGCCGTAGCGCCGATGGTAGTGTGGGGTCTCCCCATG<br>CGAGAGTAGGGAACTGCCAGGCATCAAATAAAACGAAAGGCTCAGTCGAA<br>AGACTGGGCCTTTCGTTTTATCTGTTGTTTGTCGGTGAACT |

**Supplementary Table 3.** RNA inputs used in this study.

| Name | Length (nt) | Sequence |
|---|---|---|
| Input_01 | 169 | TTTTACCAAAAAGAAATTCGGACAAGCTTATTGCTCGTAAAAAAGAC<br>TGGGATCCAAAAAAAATATGGTGGTTTTGATAGTCCAACGGTAGCTT<br>ATTCAGTCCTAGTGGTTGCTAAGGTGGAAAAAGGGAAATCGAAGAA<br>GTTAAAATCCGTTAAAGAGTTACTAGGGAT |
| Input_02 | 45 | TGTGTACTCCTTTTATCATCTCACAGTTAAAGTCGCGAGAATAGG |
| Input_03 | 87 | AAAGACTGGGATCCAAAAAAAATATGGTGGTTTTGATAGTCCAACGG<br>TAGCTTATTCAGTCCTAGTGGTTGCTAAGGTGGAAAAAGGG |
| Input_04 | 251 | GAAAACAGAAGTACAGACAGGCGGATTCTCCAAGGAGTCAATTTTA<br>CCAAAAAGAAATTCGGACAAGCTTATTGCTCGTAAAAAAGACTGGG<br>ATCCAAAAAAAATATGGTGGTTTTGATAGTCCAACGGTAGCTTATTCA<br>GTCCTAGTGGTTGCTAAGGTGGAAAAAGGGAAATCGAAGAAGTTAA |

| | | |
|---|---|---|
| | | AATCCGTTAAAGAGTTACTAGGGATCACAATTATGGAAAGAAGTTC<br>CTTTGAAAAAAAATCCGATTG |
| Input_05 | 333 | GTGCGCAAAGTATTGTCCATGCCCCAAGTCAATATTGTCAAGAAAA<br>CAGAAGTACAGACAGGCGGATTCTCCAAGGAGTCAATTTTACCAAA<br>AAGAAATTCGGACAAGCTTATTGCTCGTAAAAAAGACTGGGATCCA<br>AAAAAATATGGTGGTTTTGATAGTCCAACGGTAGCTTATTCAGTCCT<br>AGTGGTTGCTAAGGTGGAAAAAGGGAAATCGAAGAAGTTAAAATCC<br>GTTAAAGAGTTACTAGGGATCACAATTATGGAAAGAAGTTCCTTTGA<br>AAAAAATCCGATTGACTTTTTAGAAGCTAAAGGATATAAGGAAGTTA<br>AAAAAGAC |
| Input_06 | 655 | TAGGCAAAGCAACCGCAAAATATTTCTTTTACTCTAATATCATGAAC<br>TTCTTCAAAACAGAAATTACACTTGCAAATGGAGAGATTCGCAAACG<br>CCCTCTAATCGAAACTAATGGGGAAACTGGAGAAATTGTCTGGGAT<br>AAAGGGCGAGATTTTGCCACAGTGCGCAAAGTATTGTCCATGCCCC<br>AAGTCAATATTGTCAAGAAAACAGAAGTACAGACAGGCGGATTCTC<br>CAAGGAGTCAATTTTACCAAAAAGAAATTCGGACAAGCTTATTGCTC<br>GTAAAAAAGACTGGGATCCAAAAAAATATGGTGGTTTTGATAGTCC<br>AACGGTAGCTTATTCAGTCCTAGTGGTTGCTAAGGTGGAAAAAGGG<br>AAATCGAAGAAGTTAAAATCCGTTAAAGAGTTACTAGGGATCACAAT<br>TATGGAAAGAAGTTCCTTTGAAAAAAAATCCGATTGACTTTTTAGAAG<br>CTAAAGGATATAAGGAAGTTAAAAAAGACTTAATCATTAAACTACCT<br>AAATATAGTCTTTTTGAGTTAGAAAACGGTCGTAAACGGATGCTGG<br>CTAGTGCCGGAGAATTACAAAAAGGAAATGAGCTGGCTCTGCCAA<br>GCAAATATGTGAATTTTTTATATTTAGCTAGTCATTATGAAAAGTTGA<br>AGGG |
| Input_07 | 169 | CCACTCAACGATGTGGGGACGCCGTTGCAACTTCGAGGACCTAAT<br>GTGACCGACCTAGATTCGTCATTGTGGGCAGAATGAAGTATTGGCA<br>GACATTGAGTGCCGAACAAGACCTGACCTAACGGTAAGAGAGTCT<br>CATAATACGTCCGGCCTCTTGCGCAGGtTATAT |
| Input_08 | 816 | ATGAGCCATATTCAACGGGAAACGTCTTGCTCCAGGCCGCGATTAA<br>ATTCCAACATGGATGCTGATTTATATGGGTATAAATGGGCTCGCGA<br>TAATGTCGGGCAATCAGGTGCGACAATCTATCGATTGTATGGGAAG<br>CCCGATGCGCCAGAGTTGTTTCTGAAACATGGCAAAGGTAGCGTT<br>GCCAATGATGTTACAGATGAGATGGTCAGACTAAACTGGCTGACGG<br>AATTTATGCCTCTTCCGACCATCAAGCATTTTATCCGTACTCCTGAT<br>GATGCATGGTTACTCACCACTGCGATCCCCGGGAAAACAGCATTCC<br>AGGTATTAGAAGAATATCCTGATTCAGGTGAAAATATTGTTGATGCG<br>CTGGCAGTGTTCCTGCGCCGGTTGCATTCGATTCCTGTTTGTAATT<br>GTCCTTTTAACAGCGATCGCGTATTTCGTCTCGCTCAGGCGCAATC<br>ACGAATGAATAACGGTTTGGTTGATGCGAGTGATTTTGATGACGAG<br>CGTAATGGCTGGCCTGTTGAACAAGTCTGGAAAGAAATGCATAAGC<br>TTTTGCCATTCTCACCGGATTCAGTCGTCACTCATGGTGATTTCTCA<br>CTTGATAACCTTATTTTTGACGAGGGGAAATTAATAGGTTGTATTGA<br>TGTTGGACGAGTCGGAATCGCAGACCGATACCAGGATCTTGCCAT<br>CCTATGGAACTGCCTCGGTGAGTTTTCTCCTTCATTACAGAAACGG<br>CTTTTTCAAAAATATGGTATTGATAATCCTGATATGAATAAATTGCAG<br>TTTCATTTGATGCTCGATGAGTTTTTCTAA |

**Supplementary Table 4.** Guide sequences used in this study.

| Name | Length (nt) | Sequence |
|---|---|---|
| Guide_01 | 164 | CGTTGACGAATCTTGGAGCTCCCGCTGCTTTTAAATATTTTGATACAACAATTGATCGTAAACGATATACGTCTACAAAAGAAGTTTTAGATGCCACTCTTATCCATCAATCCATCACTGGTCTTTATGAAACACGCATTGATTTGAGTCAGCTAGGAGGTGAC |
| Guide_02 | 164 | ACCAATACACGAACAAGCAGGTCACCTCCTAGCTGACTCAAATCAATGCGTGTTTCATAAAGACCAGTGATGGATTGATGGATAAGAGTGGCATCTAAAACTTCTTTTGTAGACGTATATCGTTTACGATCAATTGTTGTATCAAAATATTTAAAAGCAGCGGGAGCTCCAAGATTCGTCAACG |
| Guide_03 | 82 | GGACTATCAAAACCACCATATTTTTTTGGATCCCAGTCTTTTTTACGAGCAATAAGCTTGTCCGAATTTCTTTTTGGTAAAA |
| Guide_04 | 82 | ATCCCTAGTAACTCTTTAACGGATTTTAACTTCTTCGATTTCCCTTTTTCCACCTTAGCAACCACTAGGACTGAATAAGCTA |
| Guide_05 | 20 | ATCACACTCCCACTCCACCT |
| Guide_06 | 20 | CACACGCACATTCTCATACC |
| Guide_07 | 41 | GGACTATCAAAACCACCATATTTTTTTGGATCCCAGTCTTT |
| Guide_08 | 41 | CCCTTTTTCCACCTTAGCAACCACTAGGACTGAATAAGCTA |
| Guide_09 | 123 | GGACTATCAAAACCACCATATTTTTTTGGATCCCAGTCTTTTTTACGAGCAATAAGCTTGTCCGAATTTCTTTTTGGTAAAATTGACTCCTTGGAGAATCCGCCTGTCTGTACTTCTGTTTTC |
| Guide_10 | 123 | CAATCGGATTTTTTTCAAAGGAACTTCTTTCCATAATTGTGATCCCTAGTAACTCTTTAACGGATTTTAACTTCTTCGATTTCCCTTTTTCCACCTTAGCAACCACTAGGACTGAATAAGCTA |
| Guide_11 | 164 | GGACTATCAAAACCACCATATTTTTTTGGATCCCAGTCTTTTTTACGAGCAATAAGCTTGTCCGAATTTCTTTTTGGTAAAATTGACTCCTTGGAGAATCCGCCTGTCTGTACTTCTGTTTTCTTGACAATATTGACTTGGGGCATGGACAATACTTTGCGCAC |
| Guide_12 | 164 | GTCTTTTTTAACTTCCTTATATCCTTTAGCTTCTAAAAAGTCAATCGGATTTTTTTCAAAGGAACTTCTTTCCATAATTGTGATCCCTAGTAACTCTTTAACGGATTTTAACTTCTTCGATTTCCCTTTTTCCACCTTAGCAACCACTAGGACTGAATAAGCTA |
| Guide_13 | 325 | GGACTATCAAAACCACCATATTTTTTTGGATCCCAGTCTTTTTTACGAGCAATAAGCTTGTCCGAATTTCTTTTTGGTAAAATTGACTCCTTGGAGAATCCGCCTGTCTGTACTTCTGTTTTCTTGACAATATTGACTTGGGGCATGGACAATACTTTGCGCACTGTGGCAAAATCTCGCCCTTTATCCCAGACAATTTCTCCAGTTTCCCCATTAGTTTCGATTAGAGGGCGTTTGCGAATCTCTCCATTTGCAAGTGTAATTTCTGTTTTGAAGAAGTTCATGATATTAGAGTAAAAGAAATATTTTGCGGTTGCTTTGCCTA |
| Guide_14 | 325 | CCCTTCAACTTTTCATAATGACTAGCTAAATATAAAAAATTCACATATTTGCTTGGCAGAGCCAGCTCATTTCCTTTTTGTAATTCTCCGGCACTAGCCAGCATCCGTTTACGACCGTTTTCTAACTCAAAAAGACTATATTTAGGTAGTTTAATGATTAAGTCTTTTTTAACTTCCTTATATCCTTTAGCTTCTAAAAAGTCAATCGGATTTTTTTCAAAGGAACTTCTTTCCATAATTGTGATCCCTAGTAACTCTTTAACGGA |

| | | |
|---|---|---|
| | | TTTTAACTTCTTCGATTTCCCTTTTTCCACCTTAGCAACCACTAG GACTGAATAAGCTA |
| Guide_15 | 82 | TTCATTCTGCCCACAATGACGAATCTAGGTCGGTCACATTAGG TCCTCGAAGTTGCAACGGCGTCCCCACATCGTTGAGTGG |
| Guide_16 | 82 | ATATAACCTGCGCAAGAGGCCGGACGTATTATGAGACTCTCTT ACCGTTAGGTCAGGTCTTGTTCGGCACTCAATGTCTGCC |
| Guide_17 | 82 | GAGCCCATTTATACCCATATAAATCAGCATCCATGTTGGAATTT AATCGCGGCCTGGAGCAAGACGTTTCCCGTTGAATATG |
| Guide_18 | 82 | CTTTGCCATGTTTCAGAAACAACTCTGGCGCATCGGGCTTCCC ATACAATCGATAGATTGTCGCACCTGATTGCCCGACATT |

**Supplementary Table 5.** Sequence of other biological parts used in this study.

| Part | DNA sequence (5' to 3') | Protein Sequence |
|---|---|---|
| IGS_01 | GGGTCA | |
| IGS_02 | GGAGGG | |
| IGS_03 | GTCACC | |
| P1_helix_01 | TGACCT | |
| P1_helix_02 | CCCTCT | |
| RBS_01 | AGGAGGAA | |
| RBS_02 | AAAGAGGAGAAA | |
| HP14 | ACGTCGACTCTCGAGTGAGATTGTTGACGGTACCG TATTTT | |
| Toehold | ACCAATACACGAACAAGCAG | |
| RNA inhibitor | CGTTGACGAATCTTGGAGCTCCCGCTGCTTTTAAAT ATTTTGATACAACAATTGATCGTAAACGATATACGT CTACAAAAGAAGTTTTAGATGCCACTCTTATCCATC AATCCATCACTGGTCTTTATGAAACACGCATTGATT TGAGTCAGCTAGGAGGTGACCTGCTTGTTCGTGTA TTGGT | |
| Stem_01 | GGATCA | |
| Stem_02 | TGATCC | |
| P1_loop_WT | AAATAGCAATATTTACCTTT | |
| P1_loop_01 | AAATAGCAA | |
| P1_loop_02 | TATTTACCTTT | |
| P1_loop_03 | TAGTTACCTTT | |
| PJ23119 | TTGACAGCTAGCTCAGTCCTAGGTATAATACTAGT | |
| PJ23115 | TTTATAGCTAGCTCAGCCCTTGGTACAATGCTAGC | |
| Pbad | GCCGTCACTGCGTCTTTTACTGGCTCTTCTCGCTAA CCAAACCGGTAACCCCGCTTATTAAAAGCATTCTGT AACAAAGCGGGACCAAAGCCATGACAAAAACGCGT AACAAAAGTGTCTATAATCACGGCAGAAAAGTCCAC ATTGATTATTTGCACGGCGTCACACTTTGCTATGCC ATAACATTTTTATCCATAAGATTAGCGGATCCTACC TGACGCTTTTTATCGCAACTCTCTACTGTTTCTCCA TACCCGTTTTTTTGGGCTAGC | |
| T500 | CAAAGCCCGCCGAAAGGCGGGCTTTTTTTT | |

| | | |
|---|---|---|
| TL3S2P55 | CTCGGTACCAAAGACGAACAATAAGACGCTGAAAA<br>GCGTCTTTTTTCGTTTTGGTCC | |
| TTonB | CCTCCGACCGGAGGCTTTTGACT | |
| TrrnB | GAAGCTTGGGCCCGAACAAAAACTCATCTCAGAAG<br>AGGATCTGAATAGCGCCGTCGACCATCATCATCAT<br>CATCATTGAGTTTAAACGGTCTCCAGCTTGGCTGTT<br>TTGGCGGATGAGAGAAGATTTTCAGCCTGATACAG<br>ATTAAATCAGAACGCAGAAGCGGTCTGATAAAACA<br>GAATTTGCCTGGCGGCAGTAGCGCGGTGGTCCCA<br>CCTGACCCCATGCCGAACTCAGAAGTGAAACGCCG<br>TAGCGCCGATGGTAGTGTGGGGTCTCCCCATGCG<br>AGAGTAGGGAACTGCCAGGCATCAAATAAAACGAA<br>AGGCTCAGTCGAAAGACTGGGCCTTTCGTTTTATCT<br>GTTGTTTGTCGGTGAACT | |
| Ribozyme | AAAAGTTATCAGGCATGCACCTGGTAGCTAGTCTTT<br>AAACCAATAGATTGCATCGGTTTAAAAGGCAAGACC<br>GTCAAATTGCGGGAAAGGGGTCAACAGCCGTTCAG<br>TACCAAGTCTCAGGGGAAACTTTGAGATGGCCTTG<br>CAAAGGGTATGGTAATAAGCTGACGGACATGGTCC<br>TAACCACGCAGCCAAGTCCTAAGTCAACAGATCTT<br>CTGTTGATATGGATGCAGTTCACAGACTAAATGTCG<br>GTCGGGGAAGATGTATTCTTCTCATAAGATATAGTC<br>GGACCTCTCCTTAATGGGAGCTAGCGGATGAAGTG<br>ATGCAACACTGGAGCCGCTGGGAACTAATTTGTAT<br>GCGAAAGTATATTGATTAGTTTTGGAGTACTCG | |
| dRibozyme | AAAAGTTATCAGGCATGCACCTGGTAGCTAGTCTTT<br>AAACCAATAGATTGCATCGGTTTAAAAGGCAAGACC<br>GTCAAATTGCGGGAAAGGGGTCAACAGCCGTTCAG<br>TACCAAGTCTCAGGGGAAACTTTGAGATGGCCTTG<br>CAAAGGGTATGGTAATAAGCTGACGGACATGGTCC<br>TAACCACGCAGCCAAGTCCTAAGTCAACAGATCTT<br>CTGTTGATATGGATGCAGTTCACAAACTAAATGTCG<br>GTCGGGGAAGATGTATTCTTCTCATAAGATATAGTC<br>GGACCTCTCCTTAATGGGAGCTAGCGGATGAAGTG<br>ATGCAACACTGGAGCCGCTGGGAACTAATTTGTAT<br>GCGAAAGTATATTGATTAGTTTTGGAGTACTCG | |
| sgRNA_01 | AATTAGATGGTGATGTTAATGTTTTAGAGCTAGAAA<br>TAGCAAGTTAAAATAAGGCTAGTCCGTTATCAACTT<br>GAAAAAGTGGCACCGAGTCGGTGCTTTTTTT | |
| mCherry2 | ATGGTGAGTAAAGGAGAAGAAAACAACTTAGCTAT<br>CATTAAAGAGTTCATGCGCTTCAAAGTTCACATGGA<br>GGGTTCTGTTAACGGTCACGAGTTCGAGATCGAAG<br>GCGAAGGCGAGGGCCGTCCGTATGAAGGCACCCA<br>GACCGCCAAACTGAAAGTGACTAAAGGCGGCCCG<br>CTGCCTTTTGCGTGGGACATCCTGAGCCCGCAATT<br>TATGTACGGTTCTAAAGCGTATGTTAAACACCCAGC<br>GGATATCCCGGACTATCTGAAGCTGTCTTTTCCGG<br>AAGGTTTCAACTGGGAACGCGTAATGAATTTTGAAG<br>ATGGTGGTGTCGTGACCGTCACTCAGGACTCCTCC<br>CTTCAGGATGGCGAGTTCATCTATAAAGTTAAACTG | MVSKGEENNLAIIK<br>EFMRFKVHMEGSV<br>NGHEFEIEGEGEG<br>RPYEGTQTAKLKV<br>TKGGPLPFAWDILS<br>PQFMYGSKAYVKH<br>PADIPDYLKLSFPE<br>GFNWERVMNFED<br>GGVVTVTQDSSLQ<br>DGEFIYKVKLRGTN<br>FPSDGPVMQCRT |

| | | |
|---|---|---|
| | CGTGGTACTAATTTTCCATCTGATGGCCCGGTGAT GCAGTGTAGGACGATGGGTTGGGAGGCGTCTACC GAACGCATGTATCCGGAAGATGGTGCGCTGAAAGG CGAAATTAAACAGCGCCTGAAACTGAAAGATGGCG GCCATTATGACGCTGAAGTGAAAACCACGTACAAA GCCAAGAAACCTGTGCAGCTGCCTGGCGCGTACAA TGTGGATATTAAACTGGACATCTTATCTCATAATGA AGATTATACGATCGTAGAGCAATATGAGCGCGCGG AGGGTCGTCATTCTACCGGTGGCATGGATGAACTA TACAAATAA | MGWEASTERMYP EDGALKGEIKQRLK LKDGGHYDAEVKT TYKAKKPVQLPGA YNVDIKLDILSHNE DYTIVEQYERAEG RHSTGGMDELYK* |
| sfGFP | ATGAGCAAAGGAGAAGAACTTTTCACTGGAGTTGT CCCAATTCTTGTTGAATTAGATGGTGATGTTAATGG GCACAAATTTTCTGTCCGTGGAGAGGGTGAAGGTG ATGCTACAAACGGAAAACTCACCCTTAAATTTATTT GCACTACTGGAAAACTACCTGTTCCGTGGCCAACA CTTGTCACTACTCTGACCTATGGTGTTCAATGCTTT TCCCGTTATCCGGATCACATGAAACGGCATGACTTT TTCAAGAGTGCCATGCCCGAAGGTTATGTACAGGA ACGCACTATATCTTTCAAAGATGACGGGACCTACAA GACGCGTGCTGAAGTCAAGTTTGAAGGTGATACCC TTGTTAATCGTATCGAGTTAAAGGGTATTGATTTTAA AGAAGATGGAAACATTCTTGGACACAAACTCGAGT ACAACTTTAACTCACACAATGTATACATCACGGCAG ACAAACAAAAGAATGGAATCAAAGCTAACTTCAAAA TTCGCCACAACGTTGAAGATGGTTCCGTTCAACTA GCAGACCATTATCAACAAAATACTCCAATTGGCGAT GGCCCTGTCCTTTTACCAGACAACCATTACCTGTC GACACAATCTGTCCTTTCGAAAGATCCCAACGAAAA GCGTGACCACATGGTCCTTCTTGAGTTTGTAACTG CTGCTGGGATTACACATGGCATGGATGAGCTCTAC AAATAA | MSKGEELFTGVVPI LVELDGDVNGHKF SVRGEGEGDATNG KLTLKFICTTGKLPV PWPTLVTTLTYGV QCFSRYPDHMKRH DFFKSAMPEGYVQ ERTISFKDDGTYKT RAEVKFEGDTLVN RIELKGIDFKEDGNI LGHKLEYNFNSHN VYITADKQKNGIKA NFKIRHNVEDGSV QLADHYQQNTPIG DGPVLLPDNHYLS TQSVLSKDPNEKR DHMVLLEFVTAAGI THGMDELYK* |
| FMO | ATGGCGACCCGTATTGCAATTCTGGGCGCAGGCCC ATCGGGTATGGCGCAATTGCGTGCGTTTCAAAGCG CACAAGAGAAAGGCGCTGAGATCCCGGAGTTGGTT TGTTTTGAGAAACAGGCGGACTGGGGTGGCCAGT GGAACTATACTTGGCGTACCGGTCTGGACGAAAAC GGCGAACCGGTCCACAGCTCCATGTACCGTTACCT GTGGTCCAACGGTCCGAAAGAATGTTTGGAGTTTG CTGATTACACCTTTGATGAACACTTTGGTAAGCCAA TTGCCAGCTACCCACCGCGTGAAGTGCTGTGGGAC TATATCAAGGGTCGCGTGGAAAAGGCGGGTGTCC GCAAATACATCCGTTTCAATACCGCGGTTCGTCATG TTGAGTTCAATGAGGATTCTCAGACCTTTACTGTGA CGGTTCAGGACCATACCACTGACACCATCTATAGC GAAGAATTTGACTATGTGGTTTGCTGTACCGGTCAC TTCAGCACCCCGTATGTCCCGGAGTTCGAAGGCTT CGAAAAGTTCGGTGGTCGTATTCTGCATGCCCACG ACTTTCGTGATGCGCTGGAGTTCAAGGATAAGACC GTTCTGTTGGTGGGCAGCTCGTACTCTGCGGAAGA TATTGGCAGCCAGTGCTACAAGTATGGCGCGAAGA AACTGATTAGCTGCTATCGCACCGCACCGATGGGT | MATRIAILGAGPSG MAQLRAFQSAQEK GAEIPELVCFEKQA DWGGQWNYTWRT GLDENGEPVHSSM YRYLWSNGPKECL EFADYTFDEHFGK PIASYPPREVLWDY IKGRVEKAGVRKYI RFNTAVRHVEFNE DSQTFTVTVQDHT TDTIYSEEFDYVVC CTGHFSTPYVPEF EGFEKFGGRILHAH DFRDALEFKDKTVL LVGSSYSAEDIGSQ CYKYGAKKLISCYR TAPMGYKWPENW DERPNLVRVDTEN AYFADGSSEKVDAI |

| | | |
|---|---|---|
| | TACAAATGGCCGGAGAACTGGGACGAGCGTCCGA ACCTGGTGCGTGTGGATACCGAGAATGCTTACTTC GCAGATGGTTCTTCGGAGAAAGTTGATGCCATCAT CCTGTGCACCGGTTACATCCACCACTTCCCGTTTCT GAATGACGACTTGCGCCTGGTGACCAACAATCGCC TGTGGCCGCTGAACCTGTACAAGGGCGTTGTTTGG GAGGATAATCCGAAGTTCTTCTACATTGGTATGCAA GACCAATGGTACAGCTTCAACATGTTCGATGCCCA AGCTTGGTATGCGCGTGATGTGATCATGGGCCGTT TGCCGTTGCCGAGCAAAGAAGAAATGAAGGCCGAC AGCATGGCGTGGCGCGAGAAAGAGCTGACGCTGG TCACGGCTGAAGAGATGTATACCTACCAGGGTGAC TATATCCAGAACCTGATCGACATGACCGATTATCCG AGCTTTGATATTCCGGCGACGAACAAAACGTTCCT GGAATGGAAACATCATAAGAAAGAGAACATCATGA CGTTTCGCGACCACAGCTATCGCTCCCTGATGACC GGCACGATGGCTCCAAAGCACCATACCCCGTGGAT CGATGCTCTGGACGACAGCCTGGAGGCTTACCTGA GCGACAAGTCCGAAATCCCGGTGGCAAAAGAGGC CTGATAA | ILCTGYIHHFPFLND DLRLVTNNRLWPL NLYKGVVWEDNPK FFYIGMQDQWYSF NMFDAQAWYARD VIMGRLPLPSKEEM KADSMAWREKELT LVTAEEMYTYQGD YIQNLIDMTDYPSF DIPATNKTFLEWKH HKKENIMTFRDHS YRSLMTGTMAPKH HTPWIDALDDSLEA YLSDKSEIPVAKEA ** |
| MHT | ATGAGCACCGTGGCGAACATTGCGCCGGTGTTTAC CGGCGATTGCAAAACCATTCCGACCCCGGAAGAAT GCGCGACCTTTCTGTATAAAGTGGTGAACAGCGGC GGCTGGGAAAAATGCTGGGTGGAAGAAGTGATTCC GTGGGATCTGGGCGTGCCGACCCCGCTGGTGCTG CATCTGGTGAAAAACAACGCGCTGCCGAACGGCAA AGGCCTGGTGCCGGGCTGCGGCGGCGGCTATGAT GTGGTGGCGATGGCGAACCCGGAACGCTTTATGG TGGGCCTGGATATTAGCGAAAACGCGCTGAAAAAA GCGCGCGAAACCTTTAGCACCATGCCGAACAGCAG CTGCTTTAGCTTTGTGAAAGAAGATGTGTTTACCTG GCGCCCGGAACAGCCGTTTGATTTTATTTTTGATTA TGTGTTTTTTTGCGCGATTGATCCGAAAATGCGCCC GGCGTGGGGCAAAGCGTATGAACTGCTGAAACCG GATGGCGAACTGATTACCCTGATGTATCCGATTAC CAACCATGAAGGCGGCCCGCCGTTTAGCGTGAGC GAAAGCGAATATGAAAAAGTGCTGGTGCCGCTGGG CTTTAAACAGCTGAGCCTGGAAGATTATAGCGATCT GGCGGTGGAACCGCGCAAAGGCAAAGAAAAACTG GCGCGCTGGAAAAAAATGAACAACTGA | MSTVANIAPVFTGD CKTIPTPEECATFL YKVVNSGGWEKC WVEEVIPWDLGVP TPLVLHLVKNNALP NGKGLVPGCGGG YDVVAMANPERFM VGLDISENALKKAR ETFSTMPNSSCFS FVKEDVFTWRPEQ PFDFIFDYVFFCAID PKMRPAWGKAYEL LKPDGELITLMYPIT NHEGGPPFSVSES EYEKVLVPLGFKQL SLEDYSDLAVEPR KGKEKLARWKKMN N* |
| dCas9 | ATGGACAAGAAGTATTCTATCGGACTGGCTATCGG GACTAATAGCGTCGGGTGGGCCGTGATCACTGAC GAGTACAAGGTGCCCTCTAAGAAGTTCAAGGTGCT CGGGAACACCGACCGGCATTCCATCAAGAAAAATC TGATCGGAGCTCTCCTCTTTGATTCAGGGGAAACC GCTGAAGCAACCCGCCTCAAGCGGACTGCTAGAC GGCGGTACACCAGGAGGAAGAACCGGATTTGTTAC CTTCAAGAGATATTCTCCAACGAAATGGCAAAGGTC GACGACAGCTTCTTCCATAGGCTGGAAGAATCATT CCTCGTGGAAGAGGATAAGAAGCATGAACGGCATC CCATCTTCGGTAATATCGTCGACGAGGTGGCCTAT | MDKKYSIGLAIGTN SVGWAVITDEYKV PSKKFKVLGNTDR HSIKKNLIGALLFDS GETAEATRLKRTA RRRYTRRKNRICYL QEIFSNEMAKVDD SFFHRLEESFLVEE DKKHERHPIFGNIV DEVAYHEKYPTIYH LRKKLVDSTDKADL |

| | |
|---|---|
| CACGAGAAATACCCAACCATCTACCATCTTCGCAAA | RLIYLALAHMIKFR |
| AAGCTGGTGGACTCAACCGACAAGGCAGACCTCC | GHFLIEGDLNPDNS |
| GGCTTATCTACCTGGCCCTGGCCCACATGATCAAG | DVDKLFIQLVQTYN |
| TTCAGAGGCCACTTCCTGATCGAGGGCGACCTCAA | QLFEENPINASGVD |
| TCCTGACAATAGCGATGTGGATAAACTGTTCATCCA | AKAILSARLSKSRR |
| GCTGGTGCAGACTTACAACCAGCTCTTTGAAGAGA | LENLIAQLPGEKKN |
| ACCCCATCAATGCAAGCGGAGTCGATGCCAAGGCC | GLFGNLIALSLGLT |
| ATTCTGTCAGCCCGGCTGTCAAAGAGCCGCAGACT | PNFKSNFDLAEDA |
| TGAGAATCTTATCGCTCAGCTGCCGGGTGAAAAGA | KLQLSKDTYDDDL |
| AAAATGGACTGTTCGGGAACCTGATTGCTCTTTCAC | DNLLAQIGDQYADL |
| TTGGGCTGACTCCCAATTTCAAGTCTAATTTCGACC | FLAAKNLSDAILLS |
| TGGCAGAGGATGCCAAGCTGCAACTGTCCAAGGAC | DILRVNTEITKAPLS |
| ACCTATGATGACGATCTCGACAACCTCCTGGCCCA | ASMIKRYDEHHQD |
| GATCGGTGACCAATACGCCGACCTTTTCCTTGCTG | LTLLKALVRQQLPE |
| CTAAGAATCTTTCTGACGCCATCCTGCTGTCTGACA | KYKEIFFDQSKNGY |
| TTCTCCGCGTGAACACTGAAATCACCAAGGCCCCT | AGYIDGGASQEEF |
| CTTTCAGCTTCAATGATTAAGCGGTATGATGAGCAC | YKFIKPILEKMDGT |
| CACCAGGACCTGACCCTGCTTAAGGCACTCGTCCG | EELLVKLNREDLLR |
| GCAGCAGCTTCCGGAGAAGTACAAGGAAATCTTCT | KQRTFDNGSIPHQI |
| TTGACCAGTCAAAGAATGGATACGCCGGCTACATC | HLGELHAILRRQED |
| GACGGAGGTGCCTCCCAAGAGGAATTTTATAAGTT | FYPFLKDNREKIEKI |
| TATCAAACCTATCCTTGAGAAGATGGACGGCACCG | LTFRIPYYVGPLAR |
| AAGAGCTCCTCGTGAAACTGAATCGGGAGGATCTG | GNSRFAWMTRKS |
| CTGCGGAAGCAGCGCACTTTCGACAATGGGAGCAT | EETITPWNFEEVVD |
| TCCCCACCAGATCCATCTTGGGGAGCTTCACGCCA | KGASAQSFIERMT |
| TCCTTCGGCGCCAAGAGGACTTCTACCCCTTTCTTA | NFDKNLPNEKVLP |
| AGGACAACAGGGAGAAGATTGAGAAAATTCTCACT | KHSLLYEYFTVYNE |
| TTCCGCATCCCCTACTACGTGGGACCCCTCGCCAG | LTKVKYVTEGMRK |
| AGGAAATAGCCGGTTTGCTTGGATGACCAGAAAGT | PAFLSGEQKKAIVD |
| CAGAAGAAACTATCACTCCCTGGAACTTCGAAGAG | LLFKTNRKVTVKQL |
| GTGGTGGACAAGGGAGCCAGCGCTCAGTCATTCAT | KEDYFKKIECFDSV |
| CGAACGGATGACTAACTTCGATAAGAACCTCCCCA | EISGVEDRFNASLG |
| ATGAGAAGGTCCTGCCGAAACATTCCCTGCTCTAC | TYHDLLKIIKDKDFL |
| GAGTACTTTACCGTGTACAACGAGCTGACCAAGGT | DNEENEDILEDIVLT |
| GAAATATGTCACCGAAGGGATGAGGAAGCCCGCAT | LTLFEDREMIEERL |
| TCCTGTCAGGCGAACAAAAGAAGGCAATTGTGGAC | KTYAHLFDDKVMK |
| CTTCTGTTCAAGACCAATAGAAAGGTGACCGTGAA | QLKRRRYTGWGRL |
| GCAGCTGAAGGAGGACTATTTCAAGAAAATTGAAT | SRKLINGIRDKQSG |
| GCTTCGACTCTGTGGAGATTAGCGGGGTCGAAGAT | KTILDFLKSDGFAN |
| CGGTTCAACGCAAGCCTGGGTACTTACCATGATCT | RNFMQLIHDDSLTF |
| GCTTAAGATCATCAAGGACAAGGATTTTCTGGACAA | KEDIQKAQVSGQG |
| TGAGGAGAACGAGGACATCCTTGAGGACATTGTCC | DSLHEHIANLAGSP |
| TGACTCTCACTCTGTTCGAGGACCGGGAAATGATC | AIKKGILQTVKVVD |
| GAGGAGAGGCTTAAGACCTACGCCCATCTGTTCGA | ELVKVMGRHKPEN |
| CGATAAAGTGATGAAGCAACTTAAACGGAGAAGAT | IVIEMARENQTTQK |
| ATACCGGATGGGGACGCCTTAGCCGCAAACTCATC | GQKNSRERMKRIE |
| AACGGAATCCGGGACAAACAGAGCGGAAAGACCAT | EGIKELGSQILKEH |
| TCTTGATTTCCTTAAGAGCGACGGATTCGCTAATCG | PVENTQLQNEKLY |
| CAACTTCATGCAACTTATCCATGATGATTCCCTGAC | LYYLQNGRDMYVD |
| CTTTAAGGAGGACATCCAGAAGGCCCAAGTGTCTG | QELDINRLSDYDVD |
| GACAAGGTGACTCACTGCACGAGCATATCGCAAAT | AIVPQSFLADDSID |

| | | |
|---|---|---|
| | CTGGCTGGTTCACCCGCTATTAAGAAGGGTATTCT | NKVLTRSDKNRGK |
| | CCAGACCGTGAAAGTCGTGGACGAGCTGGTCAAG | SDNVPSEEVVKKM |
| | GTGATGGGTCGCCATAAACCAGAGAACATTGTCAT | KNYWRQLLNAKLIT |
| | CGAGATGGCCAGGGAAAACCAGACTACCCAGAAG | QRKFDNLTKAERG |
| | GGACAGAAGAACAGCAGGGAGCGGATGAAAAGAA | GLSELDKAGFIKRQ |
| | TTGAGGAAGGGATTAAGGAGCTCGGGTCACAGATC | LVETRQITKHVAQIL |
| | CTTAAAGAGCACCCGGTGGAAAACACCCAGCTTCA | DSRMNTKYDENDK |
| | GAATGAGAAGCTCTATCTGTACTACCTTCAAAATGG | LIREVKVITLKSKLV |
| | ACGCGATATGTATGTGGACCAAGAGCTTGATATCA | SDFRKDFQFYKVR |
| | ACAGGCTCTCAGACTACGACGTGGACGCCATCGTC | EINNYHHAHDAYLN |
| | CCTCAGAGCTTCCTCGCAGACGACTCAATTGACAA | AVVGTALIKKYPAL |
| | TAAGGTGCTGACTCGCTCAGACAAGAACCGGGGAA | ESEFVYGDYKVYD |
| | AGTCAGATAACGTGCCCTCAGAGGAAGTCGTGAAA | VRKMIAKSEQEIGK |
| | AAGATGAAGAACTATTGGCGCCAGCTTCTGAACGC | ATAKYFFYSNIMNF |
| | AAAGCTGATCACTCAGCGGAAGTTCGACAATCTCA | FKTEITLANGEIRKA |
| | CTAAGGCTGAGAGGGGCGGACTGAGCGAACTGGA | PLIETNGETGEIVW |
| | CAAAGCAGGATTCATTAAACGGCAACTTGTGGAGA | DKGRDFATVRKVL |
| | CTCGGCAGATTACTAAACATGTCGCCCAAATCCTTG | SMPQVNIVKKTEV |
| | ACTCACGCATGAATACCAAGTACGACGAAAACGAC | QTGGFSKESILPKR |
| | AAACTTATCCGCGAGGTGAAGGTGATTACCCTGAA | NSDKLIARKKDWD |
| | GTCCAAGCTGGTCAGCGATTTCAGAAAGGACTTTC | PKKYGGFDSPTVA |
| | AATTCTACAAAGTGCGGGAGATCAATAACTATCATC | YSVLVVAKVEKGK |
| | ATGCTCATGACGCATATCTGAATGCCGTGGTGGGA | SKKLKSVKELLGITI |
| | ACCGCCCTGATCAAGAAGTACCCAGCACTGGAAAG | MERSSFEKNPIDFL |
| | CGAGTTCGTGTACGGAGACTACAAGGTCTACGACG | EAKGYKEVKKDLII |
| | TGCGCAAGATGATTGCCAAATCTGAGCAGGAGATC | KLPKYSLFELENGR |
| | GGAAAGGCCACCGCAAAGTACTTCTTCTACAGCAA | KRMLASAGELQKG |
| | CATCATGAATTTCTTCAAGACCGAAATCACCCTTGC | NELALPSKYVNFLY |
| | AAACGGTGAGATCCGGAAGGCGCCGCTCATCGAG | LASHYEKLKGSPE |
| | ACTAATGGGGAGACTGGCGAAATCGTGTGGGACAA | DNEQKQLFVEQHK |
| | GGGCAGAGATTTCGCTACCGTGCGCAAAGTGCTTT | HYLDEIIEQISEFSK |
| | CTATGCCTCAAGTGAACATCGTGAAGAAAACCGAG | RVILADANLDKVLS |
| | GTGCAAACCGGAGGCTTTTCTAAGGAATCAATCCT | AYNKHRDKPIREQ |
| | CCCCAAGCGCAACTCCGACAAGCTCATTGCAAGGA | AENIIHLFTLTNLGA |
| | AGAAGGATTGGGACCCTAAGAAGTACGGCGGATTC | PAAFKYFDTTIDRK |
| | GATTCACCAACTGTGGCTTATTCTGTCCTGGTCGTG | RYTSTKEVLDATLI |
| | GCTAAGGTGGAAAAAGGAAAGTCTAAGAAGCTCAA | HQSITGLYETRIDL |
| | GAGCGTGAAGGAACTGCTGGGTATCACCATTATGG | SQLGGD* |
| | AGCGCAGCTCCTTCGAGAAGAACCCAATTGACTTT | |
| | CTCGAAGCCAAAGGTTACAAGGAAGTCAAGAAGGA | |
| | CCTTATCATCAAGCTCCCAAAGTATAGCCTGTTCGA | |
| | ACTGGAGAATGGGCGGAAGCGGATGCTCGCCTCC | |
| | GCTGGCGAACTTCAGAAGGGTAATGAGCTGGCTCT | |
| | CCCCTCCAAGTACGTGAATTTCCTCTACCTTGCAAG | |
| | CCATTACGAGAAGCTGAAGGGGAGCCCCGAGGAC | |
| | AACGAGCAAAAGCAACTGTTTGTGGAGCAGCATAA | |
| | GCATTATCTGGACGAGATCATTGAGCAGATTTCCG | |
| | AGTTTTCTAAACGCGTCATTCTCGCTGATGCCAACC | |
| | TCGATAAAGTCCTTAGCGCATACAATAAGCACAGA | |
| | GACAAACCAATTCGGGAGCAGGCTGAGAATATCAT | |
| | CCACCTGTTCACCCTCACCAATCTTGGTGCCCCTG | |

| | |
|---|---|
| | CCGCATTCAAGTACTTCGACACCACCATCGACCGG<br>AAACGCTATACCTCCACCAAAGAAGTGCTGGACGC<br>CACCCTCATCCACCAGAGCATCACCGGACTTTACG<br>AAACTCGGATTGACCTCTCACAGCTCGGAGGGGAT<br>TGA |

**Supplementary Table 6.** Strains used in this study.

| Strain Name | Genotype | Figure |
|---|---|---|
| MG1655 | $F^-$ $\lambda^-$ $ilvG^-$ rfb-50 rph-1 | F2, F4, F5 |
| TG1 | K-12 *supE thi-1 Δ(lac-proAB) Δ(mcrB-hsdSM)5,* ($r_K^- m_K^-$) F' [*traD36 proAB*$^+$ *lacI*$^q$ *lacZΔM15*] | F1, F3 |

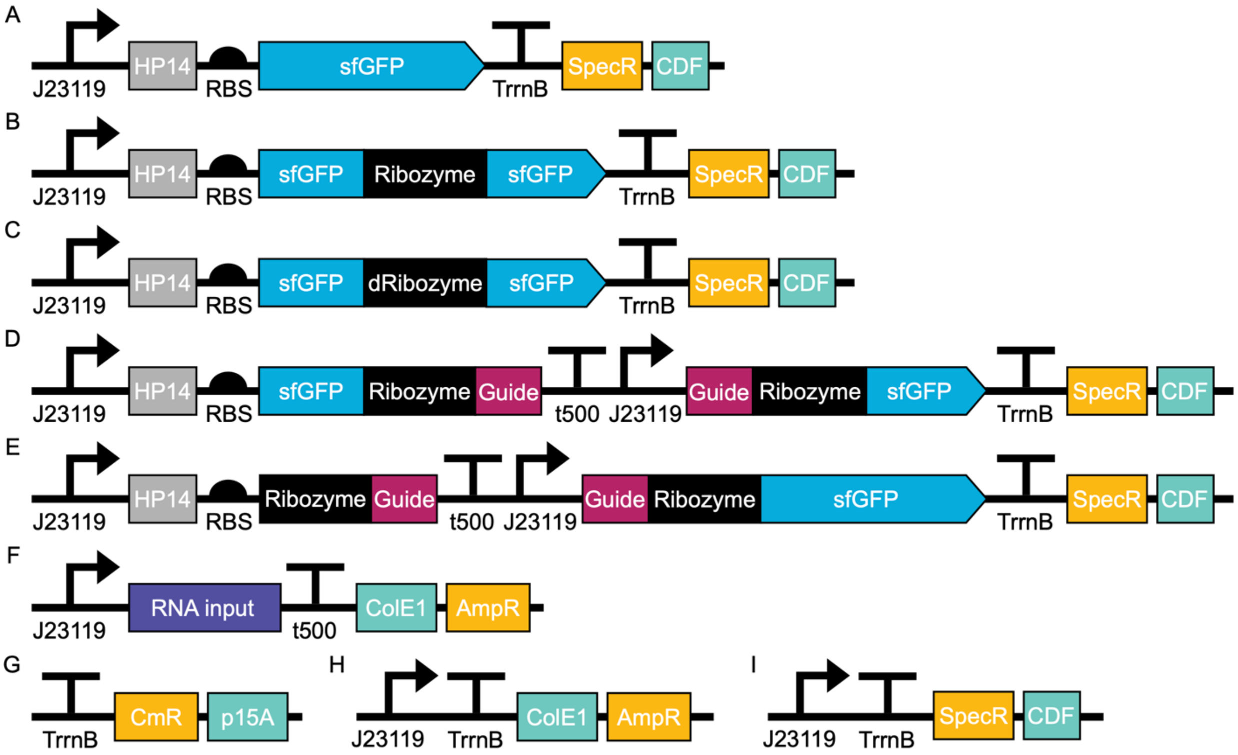

**Supplementary Fig. 1. Schematic of representative DNA plasmids used in this study**. (**A**) sfGFP expressing plasmid, (**B**) plasmid expressing the splicing ribozyme inserted within sfGFP, (**C**) plasmid expressing the catalytically dead splicing ribozyme within sfGFP, (**D**) plasmid expressing the split-output, RENDR-GFP system, (**E**) plasmid expressing the modular RENDR-GFP system, (**F**) plasmid expressing the RNA input, (**G**) Empty control plasmid for CmR p15A vectors, (**H**) Empty control plasmid for AmpR ColE1 vectors, (**I**) Empty control plasmid for SpecR CDF vectors.

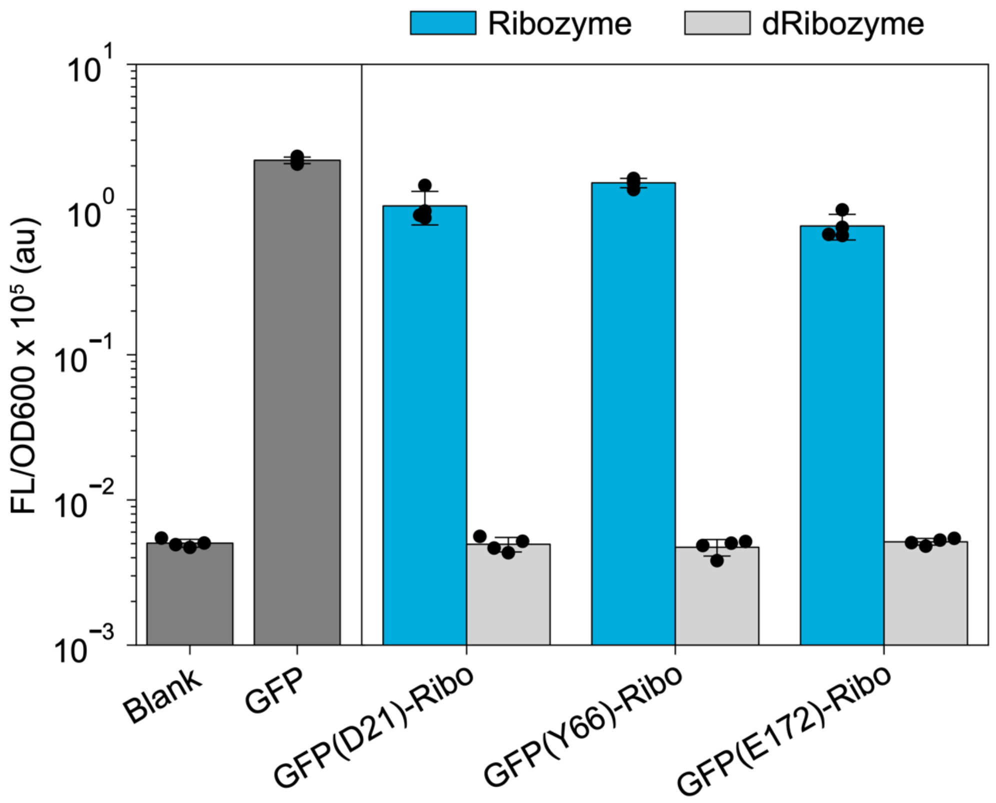

**Supplementary Fig. 2. Identifying functional ribozyme insertion sites in sfGFP.** Fluorescence characterization of ribozyme-inserted sfGFP variants (measured in units of fluorescence [FL]/optical density [OD] at 600 nm) was performed with *E. coli* transformed with an empty plasmid (Blank), a constitutively expressed sfGFP plasmid (GFP), and plasmids encoding the splicing ribozyme inserted within sfGFP at amino acid positions: D21 (GFP(D21)-Ribo), Y66 (GFP(Y66)-Ribo), and E172 (GFP(E172)-Ribo). Two variants of the ribozyme were used: a catalytically-active ribozyme (Ribozyme) and a catalytically-dead G264A mutant ribozyme (dRibozyme). Bars represent the mean values and error bars represent s.d. of $n$=4 biological replicates.

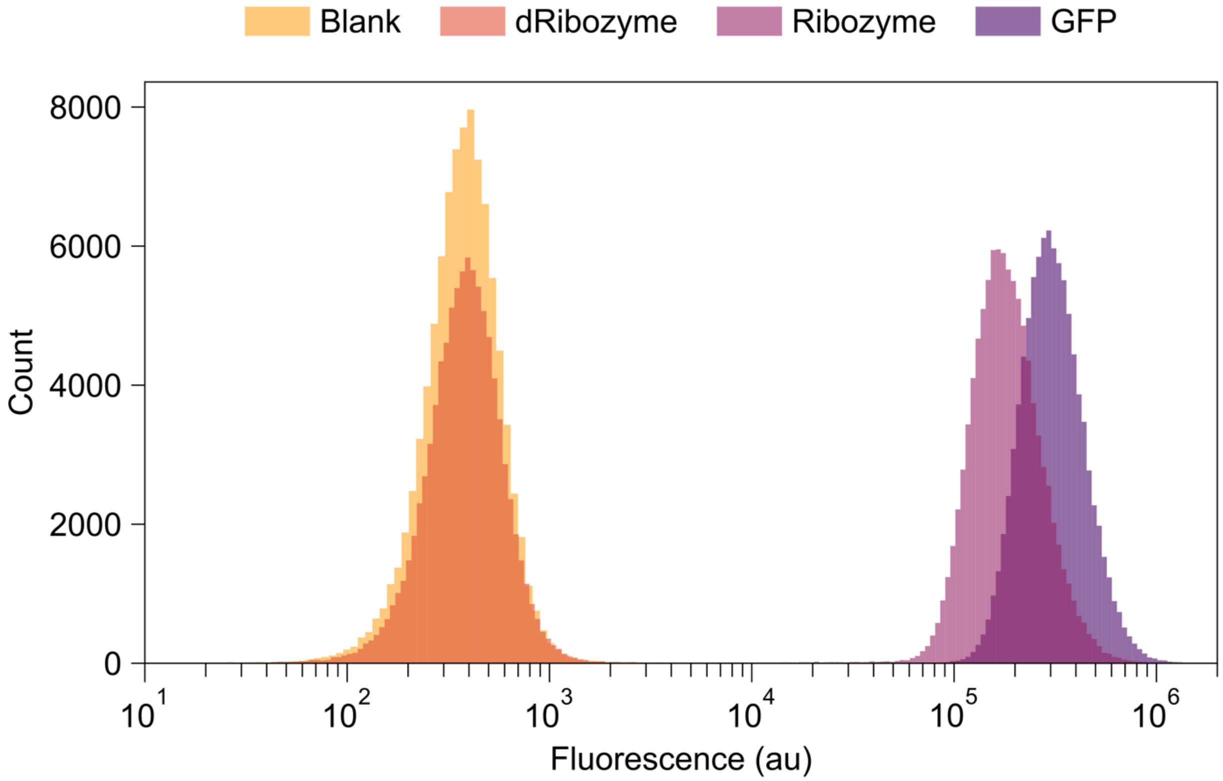

**Supplementary Fig. 3. Single cell fluorescence characterization of fluorescence-based splicing assay.** Single cell sfGFP fluorescence values of blank *E. coli* transformed with an empty control plasmid (Blank), a catalytically dead G264A mutant splicing ribozyme inserted after the first nucleotide of amino acid Y66 of sfGFP (dRibozyme), a catalytically active splicing ribozyme inserted after the first nucleotide of amino acid Y66 of GFP (Ribozyme), and an sfGFP positive control (GFP). Values measured in units of arbitrary fluorescence (au). Data shows a histogram of $n$ = 3 biological replicates.

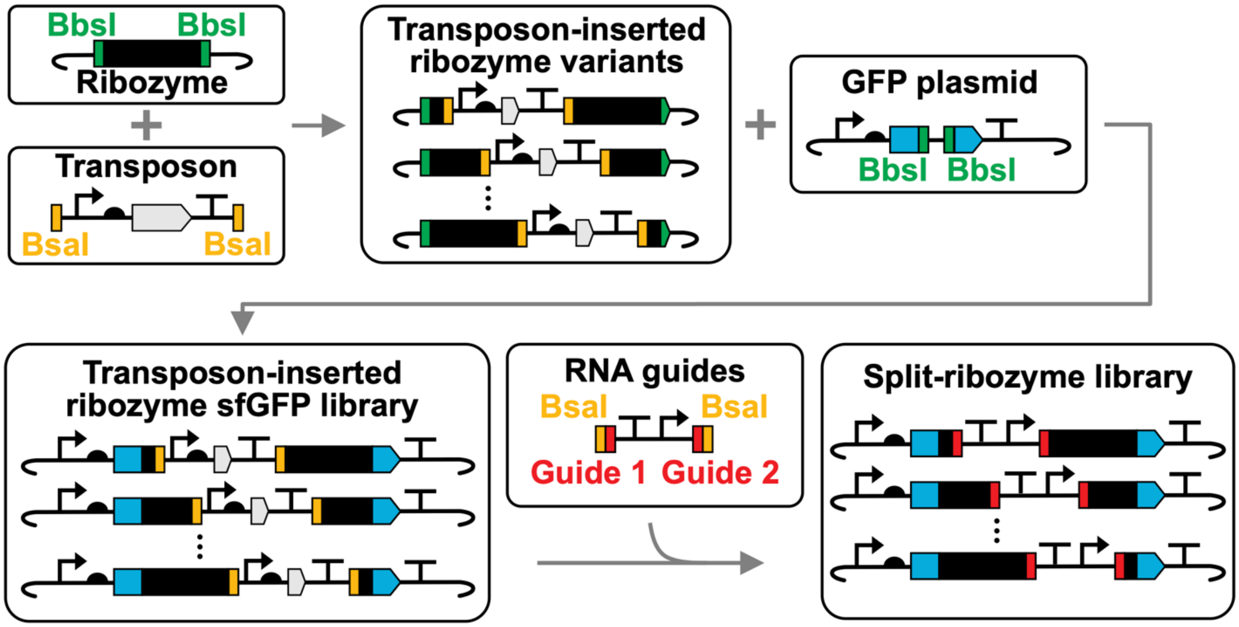

**Supplementary Fig. 4. Schematic of split-ribozyme library generation using transposon mutagenesis.** To perform *in vitro* transposon mutagenesis a synthetic transposon containing a kanamycin resistance cassette was isolated through restriction digestion and purification, and then randomly inserted into a ribozyme plasmid using Mu transposase. The subsequent transposon-inserted ribozyme variants were digested, gel purified, and cloned into a plasmid containing sfGFP. Finally, the synthetic transposon was replaced with a cassette containing RNA guide sequences, yielding the split-ribozyme library.

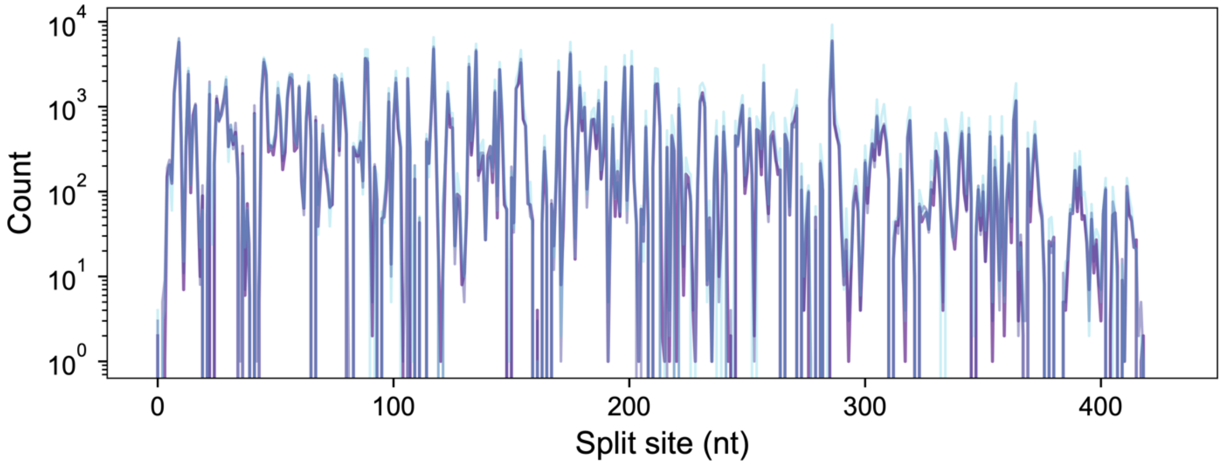

**Supplementary Fig. 5. Diversity of the split-ribozyme library.** Abundance of different ribozyme split sites within the split-ribozyme library. NGS was performed on the split-ribozyme library and the read counts processed to identify split sites. Read counts were measured from three technical replicates (blue, light purple, dark purple). This library had 93.6% coverage across all possible split sites, with 392 of 419 possible split sites present.

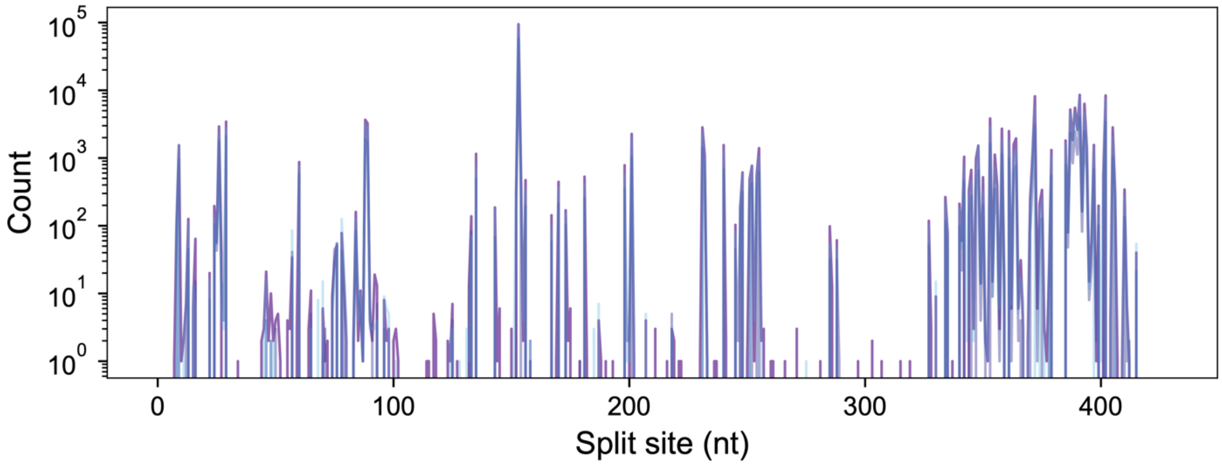

**Supplementary Fig. 6. FACS of the split-ribozyme library results in enrichment at specific split sites.** Abundance of ribozyme split sites within the split-ribozyme library following FACS. NGS was performed on the sorted split-ribozyme library and the read counts processed to identify split sites. Read counts were measured from three technical replicates (blue, light purple, dark purple). After sorting, coverage was 51.3% across all splits, with 215 of 419 possible split sites present.

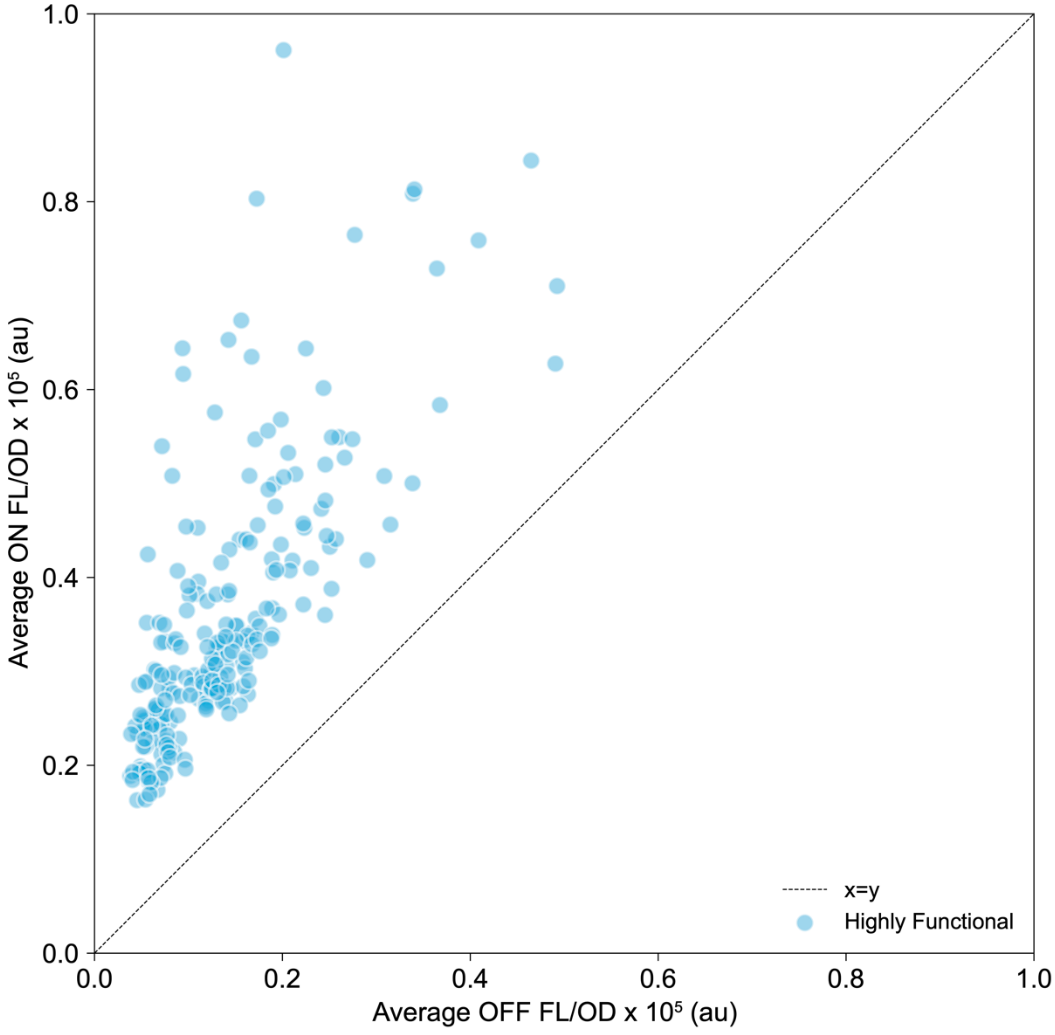

**Supplementary Fig. 7. Identification of functional split ribozymes.** Fluorescence characterization of individual split ribozyme variants from the sorted library. Fluorescence characterization (measured in units of fluorescence [FL]/optical density [OD] at 600 nm) was performed with *E. coli* transformed with plasmids encoding different split-ribozyme variants and a plasmid encoding the RNA inhibitor under the control of an AHL-inducible promoter. Fluorescence was characterized in the presence (OFF) and absence (ON) of 1 μM AHL. Functional split variants (ON>OFF) were binned based on their distance from the x=y line. Data shows mean of $n$ = 2 biological replicates.

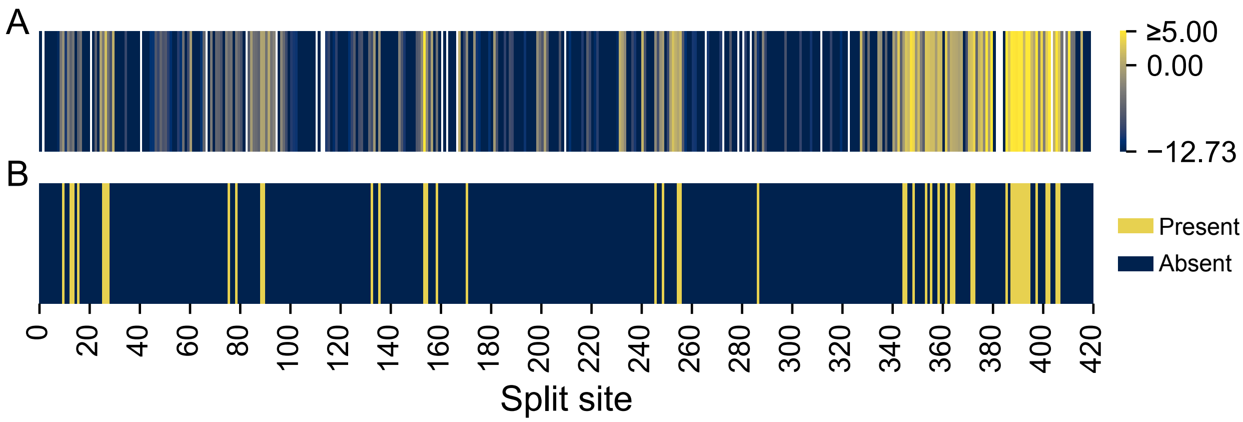

**Supplementary Fig. 8. Comparison of split variants identified through FACS-seq and individual colony screening. (A)** Average relative enrichment of split variants along the ribozyme from FACS-seq experiments (data from **Fig. 2B**). Relative enrichment values calculated from NGS of three technical replicates. **(B)** Highly functional split sites identified from individual colony screening of sorted ribozyme variants (**Supplementary Fig. 7**). Validated functional sites are yellow (Present) and those that were not present (i.e., not picked as colonies or not identified as highly functional) are blue (Absent).

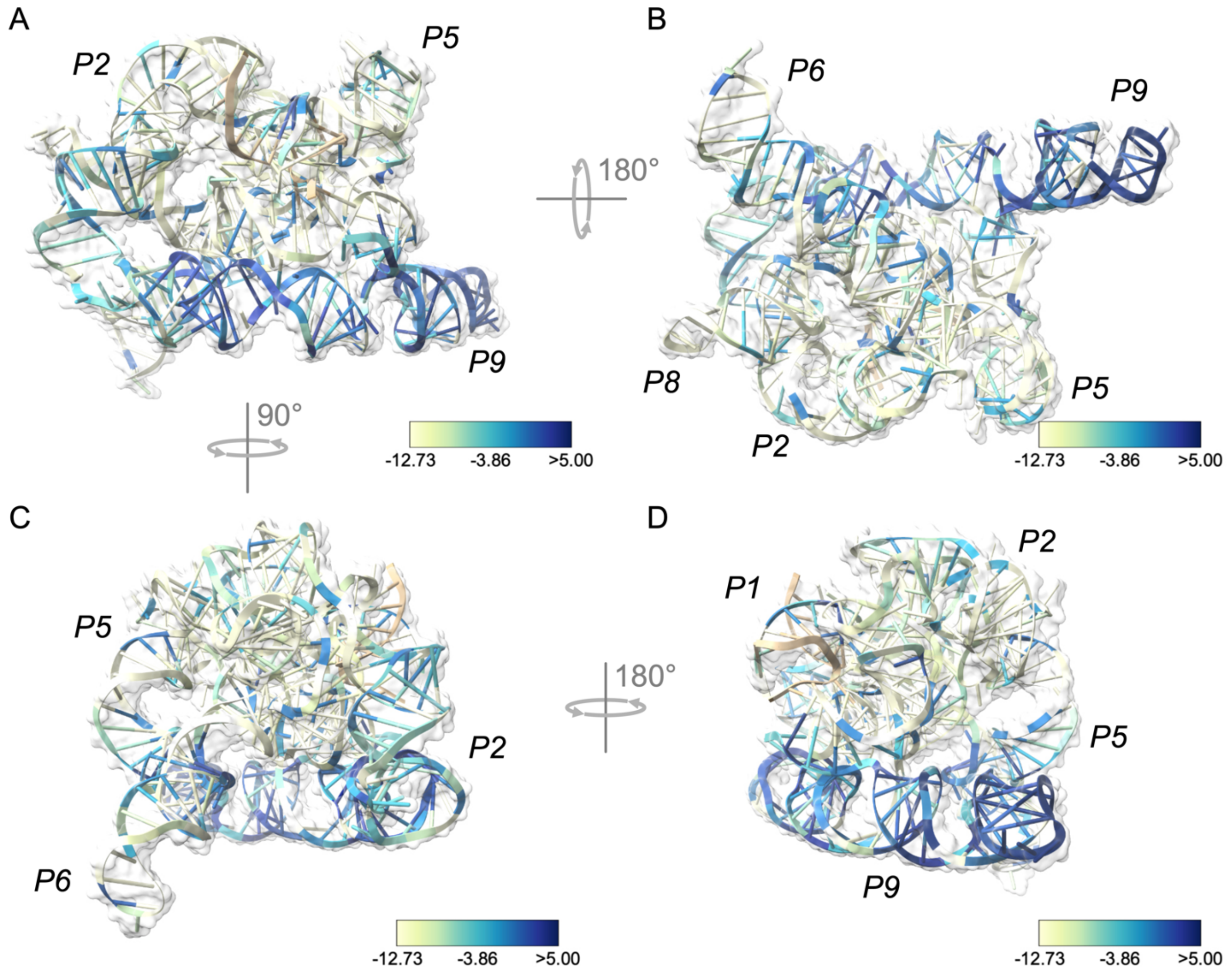

**Supplementary Fig. 9. Three-dimensional structure shows functional ribozyme split sites in surface accessible regions.** Relative enrichment of each split site overlayed onto the tertiary structure of the *Tetrahymena thermophila* ribozyme (PDB: 7EZ2)[1], at angles showing (**A**) the P9 domain wrapping around the structure, (**B**) the structure in panel (A) turned 180° along the x-axis with P2, P5, P6, P8, and P9 domains annotated (**C**) the structure in panel (A) turned 90° counterclockwise along the y-axis with the P2, P5, and P6 domains annotated, and (**D**) the structure in panel (C) turned 180° along the y-axis with the P1, P2, P5, and P9 domains annotated. Beige strands are the substrate sequences. Split sites not present in the initial library are colored in white.

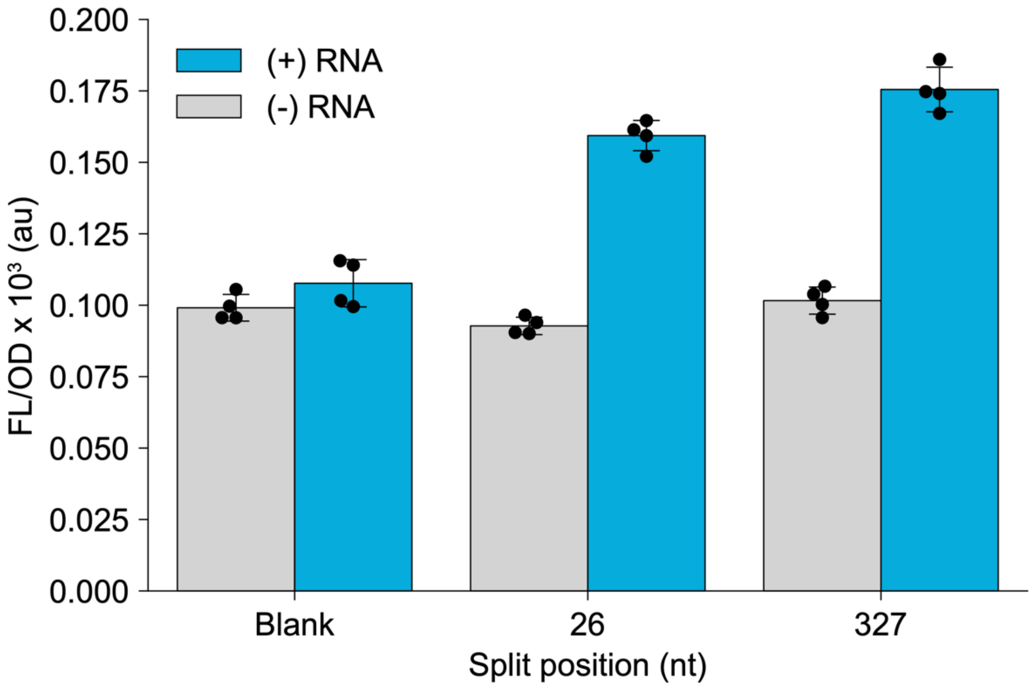

**Supplementary Fig. 10. Two enriched split sites resulted in RENDR variants with low-fold activation.** RENDR variants with split site 26 and 327 showed low (<2-fold) activation of sfGFP expression in the presence of a constitutively expressed RNA input (blue, (+) RNA) relative to these RENDR variants without RNA input (grey, (-) RNA). Fluorescence characterization (measured in units of fluorescence [FL]/optical density [OD] at 600 nm) was performed on *E. coli* co-transformed with the RENDR plasmid and either an empty plasmid or a plasmid constitutively expressing the RNA input. Autofluorescence of *E. coli* was determined using cells transformed with empty plasmids (blank). Bars show mean values and error bars represent s.d. of $n = 4$ biological replicates shown as points.

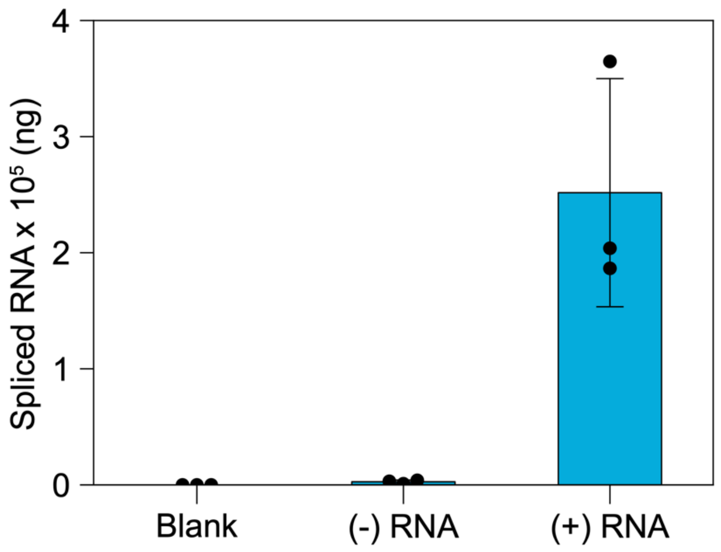

**Supplementary Fig. 11. Reverse transcription quantitative PCR (RT-qPCR) of RENDR spliced output in the presence and absence of RNA input.** Spliced RNA products from RENDR were measured using RT-qPCR using relative normalization to a standard curve (ng). Measurements were performed on total RNA extracted from *E. coli* co-transformed with the RENDR plasmid and either an empty plasmid ((-) RNA) or a plasmid constitutively expressing the RNA input ((+) RNA). *E. coli* cells transformed with empty plasmids (blank) were used as a negative control. Bars show the mean of $n$ = 3 biological replicates, of which each was measured in triplicate, shown as points with the s.d. shown as error bars.

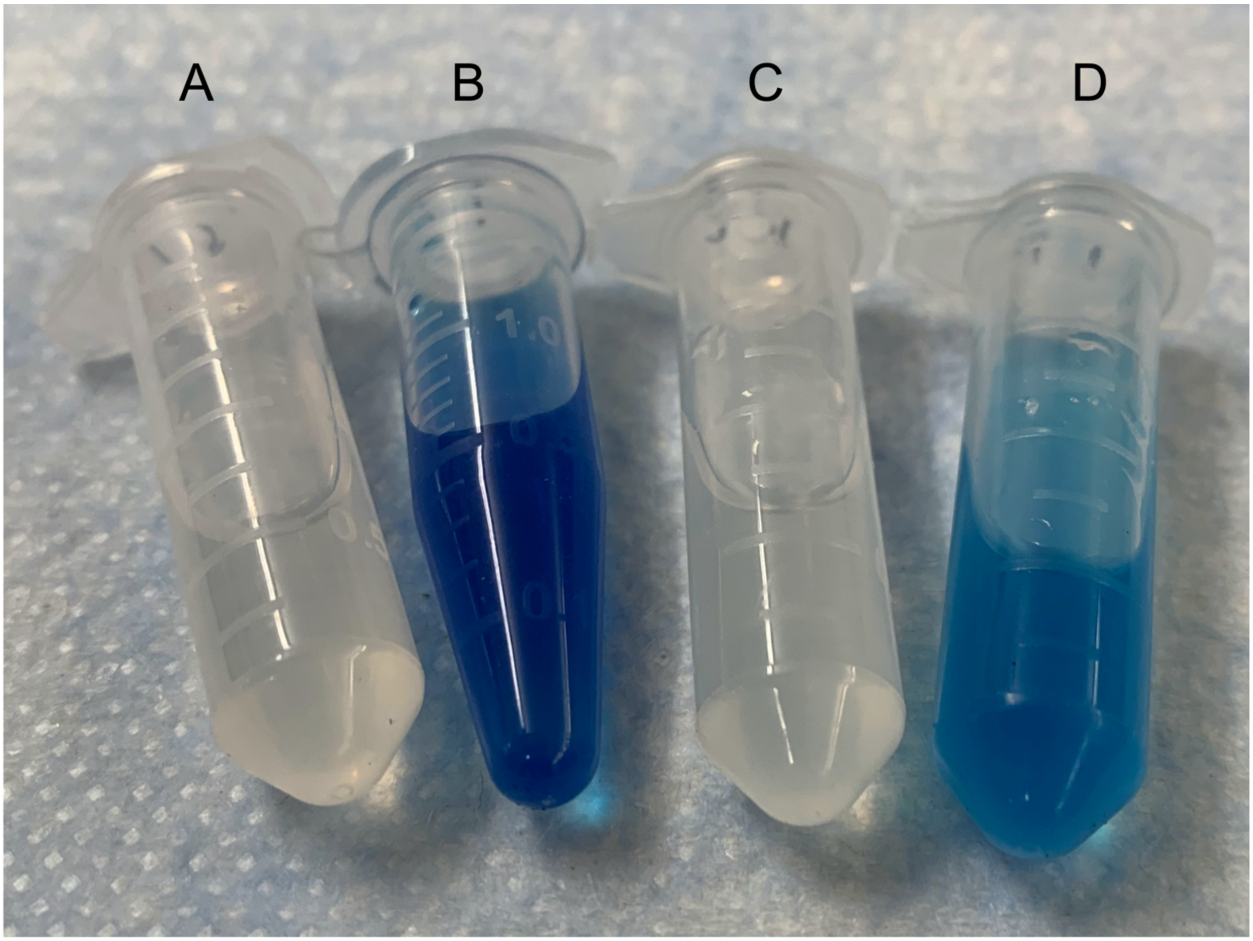

**Supplementary Fig. 12. RENDR variant using a flavin-containing monooxygenase (FMO) output produces visually discernable indigo levels upon detection of RNA.** Photograph of DMSO-extracted indigo obtained from *E. coli* cultures transformed with (**A**) an empty plasmid, (**B**) a constitutively expressed FMO protein, (**C**) a RENDR-FMO plasmid with an empty plasmid, and (**D**) a RENDR-FMO plasmid with a plasmid expressing the RNA input.

**Supplementary Note 1. Describing RENDR design principles with a thermodynamic model.**

Our goal was to establish design rules for the RENDR input detection reaction by linking the sequences of the RNA guide and RNA input to the observed output levels. We began with the consideration that the nascent ribozyme sequence has been proposed to pass through folding intermediates that then diverge into multiple folding pathways. Some fraction of intermediates rapidly fold into the native active fold, while another fraction fold into long-lived misfolded structures that are catalytically inactive. The commitment to folding into the active or inactive fraction has been shown to occur late in the folding pathway, at a point when much of the secondary and tertiary structure has formed[2].

As disruption of secondary structure elements (e.g., P1 hairpin) is known to result in misfolding of the ribozyme[3], we reasoned that by splitting the ribozyme we are promoting formation of the misfolded state from a late-folding intermediate. However, in the presence of the RNA input, interactions between the RNA guides attached to each half of the ribozyme and the RNA input form, promoting correct folding of the ribozyme into the native fold at the point of commitment. Since the RNA guides are fully complementary to the RNA input, theoretically an extended duplex can form between these species regardless of their length. However, we hypothesized that on timescales of RNA folding, which have been shown to occur on the fast timescales of transcription[4,5], only a partial intermediate "seed complex" (SC) is able to form before the folding decision is made and this SC is sufficient for promoting correct folding. Similar SC mechanisms have been proposed for gene regulatory systems involving RNA:RNA interactions that operate within kinetic windows on the timescales of transcription[6,7]. This model also captures the trend we observed in our experimental data (**Fig. 3A**) in which increasing length of interaction increases splicing, but only up to a certain length, after which it decreases.

Under these assumptions, we predicted that the rate of splicing is directly related to the rate of SC formation, which we reasoned could be captured in a sequence-function thermodynamic model.

To begin with, we assume the rate of splicing ($k_s$) is proportional to the expression of our sfGFP output, which in turn, is proportional to the observed fluorescence/optical density (FL/OD) measurements:

$$k_s \propto FL/OD \ (1)$$

The rate of splicing ($k_s$) can be estimated from the activation energy barrier ($E_a$) for the transition of the initial states (IS) to the SC using the Arrhenius equation: ($k_B$ = Boltzman's constant, T is temperature).

$$k_s \sim e^{E_a/k_BT} \ (2)$$

To estimate $E_a$, we can use a linear free energy relationship in which $E_a \sim \Delta G_{SC} - \Delta G_{IS}$ [8]:

$$E_a \sim \Delta G_{SC} - \Delta G_{IS} \ (3)$$

Taking the following definition of $\Delta G_{IS}$:

$$\Delta G_{IS} \sim \Delta G_{Guide\ 1} + \Delta G_{Guide\ 2} + \Delta G_{RNA\ input} \ (4)$$

We have:

$$FL/OD \sim k_s \sim e^{(\Delta G_{SC} - \Delta G_{Guide\ 1} + \Delta G_{Guide\ 2} + \Delta G_{RNA\ input})/k_BT} \ (5)$$

Which can be simplified to:

$$ln(FL/OD) \sim (\Delta G_{SC} - \Delta G_{Guide\ 1} + \Delta G_{Guide\ 2} + \Delta G_{RNA\ input})\ (6)$$

Therefore, using this model we predict a linear relationship between the ln(FL/OD) and the difference in free energies between the IS of the RNA guides and RNA input, and the seed complex (SC). Importantly, this model captures how intramolecular folding within each of the individual RNA species competes for folding of the SC (intermolecular interactions) which is observed in our data (**Fig. 3A**).

To test this model, we first calculated the individual ΔG terms. We estimated each ΔG term as follows:

- $\Delta G_{Guide1}$ and $\Delta G_{Guide2}$ = The minimum free energy (MFE) of the two RNA guide sequences attached to each fragment of the RENDR ribozyme. The MFE was calculated in NUPACK[9,10] with temperature = 37 °C, material = 'rna95' free energy parameter set, ensemble = 'stacking', sodium = 0.25 M, and magnesium = 0.02 M[11].
- $\Delta G_{RNAinput}$ = The MFE of the RNA input was calculated from the RNA input (not-including the terminator hairpin). The MFE was calculated in NUPACK with temperature = 37 °C, material = 'rna95' free energy parameter set, ensemble = 'stacking', sodium = 0.25 M, and magnesium = 0.02 M.
- $\Delta G_{SC}$ = The duplex binding energy of the hypothesized SC, discussed below. The MFE of the seed component of all three strands (Guide 1, Guide 2, and RNA input) was calculated in NUPACK with temperature = 37 °C, material = 'rna95' free energy parameter set, ensemble = 'stacking', sodium = 0.25 M, and magnesium = 0.02 M.

To choose the length of the sequence to set as our SC we studied how length of interaction between the RNA guide and RNA input affected overall splicing efficiency (**Fig. 3A**). From this data, it is clear that a tradeoff is observed between the competing effects of increasing the potential interaction between the RNA guides and the input RNA, and the increasing potential for interfering intramolecular folding. Studying the data, it is clear that beyond 169 bp of interaction a decrease in expression is observed, suggesting this length represents the approximate length of our proposed SC. To further support this, when we compared our fluorescent data to the predictions of eq (6) using different SC lengths increasing by increments of a single nucleotide (data not shown), we observed the strongest correlation when an SC of 173 nt was used. Using this as our SC, we see a strong correlation with an $R^2 = 0.68$ between our free energy prediction and the characterization data (**Fig. 3C**).

**Supplementary Note 2. NGS data analysis pipeline**

NGS reads processing

Forward and reverse NGS sequence reads were imported and the unique indices were used to bin reads into categories and assign directionality. For reads with at least one mutation in the index, the Hamming distance was calculated to attempt to match the mutated index with one of the indices used and thereby assign the read to the correct category. Reads that could not be conclusively assigned to an index were excluded from the dataset. Paired reads without one forward and one reverse read were also excluded. To determine the ribozyme split site, reverse reads were then aligned to reference sequences of both the ribozyme and the insertion containing the RNA guides. Briefly, we used the Biopython API[12] to perform alignments, running the Smith-Waterman local alignment algorithm with the "Nuc.4.4" substitution matrix, an open gap score of -10, and an extend gap score of -0.5. To determine where the ribozyme was split, reads were first aligned to the insertion reference sequence and reads with alignment scores below 1000 were removed along with any reads containing ambiguous alignment endpoints. The sequence following the insertion alignment endpoint, which we termed the ribozyme split sequence (RSS), was then aligned to the ribozyme reference sequence and resulting alignment scores were normalized to the length of the RSS. Reads with normalized RSS alignment scores below 4.5 were removed. Insertion sequences with RSS lengths too short to identify a unique split site, those without an RSS, and those with ambiguous split sites were also removed. Ribozyme split sites were determined from each RSS alignment.

Determining relative enrichment from FACS-seq

For each split site in the unsorted and sorted library NGS run, read counts were summed and a frequency was calculated by normalizing the read count of each split site to the total number of processed reads for that NGS run. The average frequency for technical triplicate NGS runs of the unsorted and sorted libraries was calculated and the relative enrichment of each split site was determined by dividing the average frequency of the sorted library by the average frequency of the unsorted library. Significant enrichment between the unsorted and sorted libraries was assessed through p-values using a homoscedastic two-sample t-Test with an alpha of 0.05 on the average frequencies at each split site.
